# Functional synapses between small cell lung cancer and glutamatergic neurons

**DOI:** 10.1101/2023.01.19.524045

**Authors:** Anna Schmitt, Vignesh Sakthivelu, Kristiano Ndoci, Gulzar A Wani, Marian Touet, Isabel Pintelon, Ilmars Kisis, Olta Ibruli, Julia Weber, Roman Maresch, Christina M Bebber, Jonas Goergens, Milica Jevtic, Franka Odenthal, Aleksandra Placzek, Alexandru A Hennrich, Karl-Klaus Conzelmann, Maike Boecker, Alena Heimsoeth, Gülce S Gülcüler, Ron D Jachimowicz, Julie George, Johannes Brägelmann, Silvia von Karstedt, Martin Peifer, Thorsten Persigehl, Holger Grüll, Martin L Sos, Jens Brüning, Guido Reifenberger, Matthias Fischer, Dirk Adriaensen, Reinhard Büttner, Inge Brouns, Roland Rad, Roman K Thomas, Matteo Bergami, Elisa Motori, Hans Christian Reinhardt, Filippo Beleggia

**Affiliations:** Department I of Internal Medicine, Faculty of Medicine and University Hospital Cologne, University of Cologne, Cologne, Germany; Institute of Virology, Faculty of Medicine and University Hospital Cologne, University of Cologne, Cologne, Germany; Department of Translational Genomics, Faculty of Medicine and University Hospital Cologne, University of Cologne, Cologne, Germany; Mildred Scheel School of Oncology Aachen Bonn Cologne Düsseldorf (MSSO ABCD), Faculty of Medicine and University Hospital Cologne, University of Cologne, Cologne, Germany; Cologne Excellence Cluster on Cellular Stress Response in Aging-Associated Diseases (CECAD), University of Cologne, Cologne, Germany; Laboratory of Cell Biology and Histology, Department of Veterinary Sciences, University of Antwerp, Antwerp, Belgium; Institute of Molecular Oncology and Functional Genomics, School of Medicine, Technical University of Munich, Munich, Germany; Max Planck Institute for Biology of Ageing, Cologne, Germany; Max von Pettenkofer-Institute Virology, Faculty of Medicine and Gene Center, Ludwig Maximilians University, Munich, Germany; Molecular Pathology, Institute of Pathology, Faculty of Medicine and University Hospital Cologne, University of Cologne, Cologne, Germany; Center for Molecular Medicine Cologne (CMMC), Faculty of Medicine and University Hospital Cologne, University of Cologne, Cologne, Germany; Department of Otorhinolaryngology, Head and Neck Surgery, University Hospital of Cologne, Cologne, Germany; Institute for Diagnostic and Interventional Radiology, Faculty of Medicine and University Hospital Cologne, University of Cologne, Cologne, Germany; Institute of Pathology, Faculty of Medicine and University Hospital Cologne, University of Cologne, Cologne, Germany; Department of Neuronal Control Metabolism, Max Planck Institute for Metabolism Research, Cologne, Germany; Institute of Neuropathology, Medical Faculty, Heinrich Heine University and University Hospital Düsseldorf, Düsseldorf, Germany; Department of Experimental Pediatric Oncology, University Children’s Hospital of Cologne, Medical Faculty, University of Cologne, Cologne, Germany; German Cancer Consortium (DKTK), partner site Heidelberg and German Cancer Research Center (DKFZ), Heidelberg, Germany; Institute of Genetics, University of Cologne, Cologne, Germany; Faculty of Medicine and University Hospital Cologne, University of Cologne, Cologne, Germany; Institute of Biochemistry, University of Cologne, Cologne, Germany; Dept. of Hematology and Stem Cell Transplantation, University Hospital Essen, Essen, Germany; West German Cancer Center, University Hospital Essen, Essen, Germany; DKTK Partner Site Essen/Düsseldorf, University Hospital Essen, Essen, Germany; Center for Molecular Biotechnology, University Hospital Essen, Essen, Germany

## Abstract

Small cell lung cancer (SCLC) is a highly aggressive type of lung cancer, characterized by rapid proliferation, early metastatic spread, clinical recurrence and high rate of mortality. Using *in vivo* insertional mutagenesis screening in conjunction with cross-species genomic and transcriptomic validation, we identified a strong and consistent signal for neuronal, synaptic, and glutamatergic signaling gene sets in murine and human SCLC. We show that SCLC cells have the ability to develop intimate contacts with neuronal glutamatergic terminals *in vitro*, in autochthonous primary lung tumors and in brain-engrafted tumors. These contacts can develop into *bona fide* synapses, allowing SCLC cells to receive glutamatergic inputs. Fitting with a potential oncogenic role of neuron-SCLC interactions, we show that SCLC cells derive a robust proliferation advantage when co-cultured with neurons. Moreover, the repression of glutamate release and the stimulation of the inhibitory glutamate receptor GRM8 displayed therapeutic efficacy in an autochthonous mouse model of SCLC. Therefore, following malignant transformation, SCLC cells appear to hijack glutamatergic signaling to sustain tumor growth, thereby exposing a novel entry route for therapeutic intervention.

## Introduction

Small cell lung cancer (SCLC) constitutes approximately 15% of all lung cancers in humans and is associated with a very poor prognosis ^1^. In the majority of cases, SCLC patients are diagnosed with metastatic extensive stage disease ^1^. The typical treatment is combination chemotherapy, consisting of a platinum salt and etoposide, with or without immune checkpoint blockade and optional prophylactic cranial irradiation ^1^. This intensive induction regimen induces clinical responses in more than 60% of the cases ^1^. However, these responses are almost always transient, resulting in a median overall survival of approximately one year for patients with extensive stage disease ^1^.

Recent comprehensive genomic profiling revealed that SCLC displays an almost universal bi-allelic loss of the tumor suppressors *TP53* and *RB1* ^2^. Next to these hallmark lesions, a number of additional genes, including the RB family members *RBL1* and *RBL2*, the MYC family members *MYC, MYCN* and *MYCL*, as well as *PTEN, NOTCH1, TP73, CREBBP* and *EP300* are recurrently altered in SCLC ^2,3^.

Given the highly recurrent co-occurring loss of *TP53* and *RB1*, SCLC was long thought of as a single disease entity. However, this monolithic view on SCLC biology has recently been challenged with the emergence of distinct molecular subtypes defined by the expression of lineage-related transcription factors in SCLC ^4^. Currently, at least four major molecular subtypes of SCLC are distinguished based on the expression levels of ASCL1 (subtype SCLC-A), NEUROD1 (subtype SCLC-N), POU2F3 (subtype SCLC-P) and YAP1 (subtype SCLC-Y) ^4^. These subtypes differ in the degree of neuroendocrine differentiation and use of metabolic pathways ^4^. With the advent of single cell transcriptome analyses, it is becoming clear that SCLC tumors display substantial intra-tumoral heterogeneity that appears to be dynamically altered in response to therapeutic pressure ^4^. To which degree the preservation of neuroendocrine and neuronal features inherited from the pulmonary neuroendocrine cell of origin impacts the trajectories of tumor development and drug response remains largely unclear.

In recent years, evidence suggesting that innervation impacts tumor initiation and plasticity has accumulated ^5–8^. The most compelling evidence for neuron-tumor cross talk originates from central nervous system (CNS) tumors. For instance, neurotransmitter and growth factor secretion into the microenvironment of gliomas upon neuronal activity has been shown to promote tumor cell proliferation ^9^. In addition, direct synaptic contacts between presynaptic neurons and postsynaptic cancer cells have been reported in gliomas. These synapses mediated membrane depolarizations of the postsynaptic glioma cells through AMPA glutamate receptors, resulting in increased cell division and enhanced tumor invasion ^5,10,11^. In contrast, no *bona fide* synapses have been described between neurons and non-neuronal cancers and the reported consequences of cancer-neuron interactions outside of the CNS are dependent on tumor type. For example, parasympathetic cholinergic peripheral nerves appear to promote metastatic spread of prostate cancer cells, whereas breast cancer metastasis appears to be repressed by these fibers ^5,12,13^.

SCLC derives from pulmonary neuroendocrine cells (PNECs), which in turn develop from lung epithelial progenitors of endodermal lineage ^14,15^. PNECs are innervated by different types of nerve fibers originating from the nodose, jugular and dorsal root ganglia ^16^, suggesting that SCLC-neuron interactions may be a key element in the formation and progression of SCLC and may provide clues for the development of novel therapeutic strategies.

Here, we conducted a large, unbiased *in vivo* insertional mutagenesis screen in a mouse model of SCLC and cross-referenced the results with re-analyses of all available genetic and expression data from human patients. Across species and technologies, we identified neuronal, synaptic and glutamate signaling gene sets as a prominent feature of SCLC. Further experiments uncovered the surprising ability of SCLC cells to form functional synaptic contacts with glutamatergic neurons. Moreover, pharmacologic inhibition of glutamate signaling, alone or in combination with chemotherapy, repressed SCLC growth *in vivo*.

## Results

### A genome-wide *in vivo piggyBac* mutagenesis screen in murine SCLC identifies selected insertions in synaptic genes

To search for genes and pathways that contribute to SCLC tumorigenesis *in vivo*, we performed a *piggyBac* insertional mutagenesis screen in the Ad-CMV-Cre-induced *<u>R</u>b1^fl/fl^;Tr<u>p</u>53^fl/fl^* (RP) model of SCLC ^17^. In this model, expression of the *piggyBac* transposase from the *Rosa26* locus in *Rosa26^<u>L</u>SL.PB^* mice (L) ^18^ is prevented by a *loxP*-flanked stop cassette (LSL) in the absence of exogenous Cre recombinase expression (**Extended Data Fig. 1a**). To allow for *in vivo* transposon mutagenesis, we crossed RPL mice with *ATP1-<u>S</u>2* (S, harboring 20 transposon copies on chromosome 10) or *ATP-<u>H</u>39* (H, harboring 80 transposon copies on chromosome 5) mice (**Extended Data Fig. 1b**) ^18^. SCLC tumors were induced by intratracheal instillation of Ad-CMV-Cre ^19^. Of note, the animals included in these experiments developed tumors that morphologically resembled human SCLC, as previously described for the RP model (**Extended Data Fig. 1c**) ^17,19^. We isolated and sequenced genomic DNA from 100 tumors derived from 14 untreated mice, 113 tumors from 24 mice treated with cisplatin and etoposide and 90 tumors from 20 mice treated with the anti-PD1 antibody RPM1-14 (**Extended Data Fig. 1d**).

Initial analysis revealed no significant differences in the insertion patterns of untreated and cisplatin/etoposide- or anti-PD1 antibody-treated tumors. Therefore, we pooled all the samples (303 tumors from 58 mice) for the overall statistical analysis. The significantly targeted genes in our *piggyBac* screen are distributed essentially across the entire murine genome (**Fig. 1a, Extended Data Table 1**). Of note, our screen confirms the role of genes which have been reported in SCLC, such as the tumor suppressors *Crebbp* (**Extended Data Fig. 2a**) ^20^ and *Pten* (**Extended Data Fig. 2b**) ^21^, and the transcription factors *Nfib* (**Extended Data Fig. 2c**) ^22^ and *Trp73* (**Extended Data Fig. 2d**) ^2^. Furthermore, the integration pattern in *Trp73* bears a striking resemblance with the genomic rearrangements reported in human SCLC. The transposons affect almost exclusively the long isoform, which includes the transactivation domain (**Extended Data Fig. 2d**). Similarly, the rearrangements identified in human samples lead to the expression of short transcripts (p73Δex2 and p73Δex2/3) which lack a functional transactivation domain and have dominant negative effects on the longer wildtype p73 ^2,23^.

**Figure 1:**
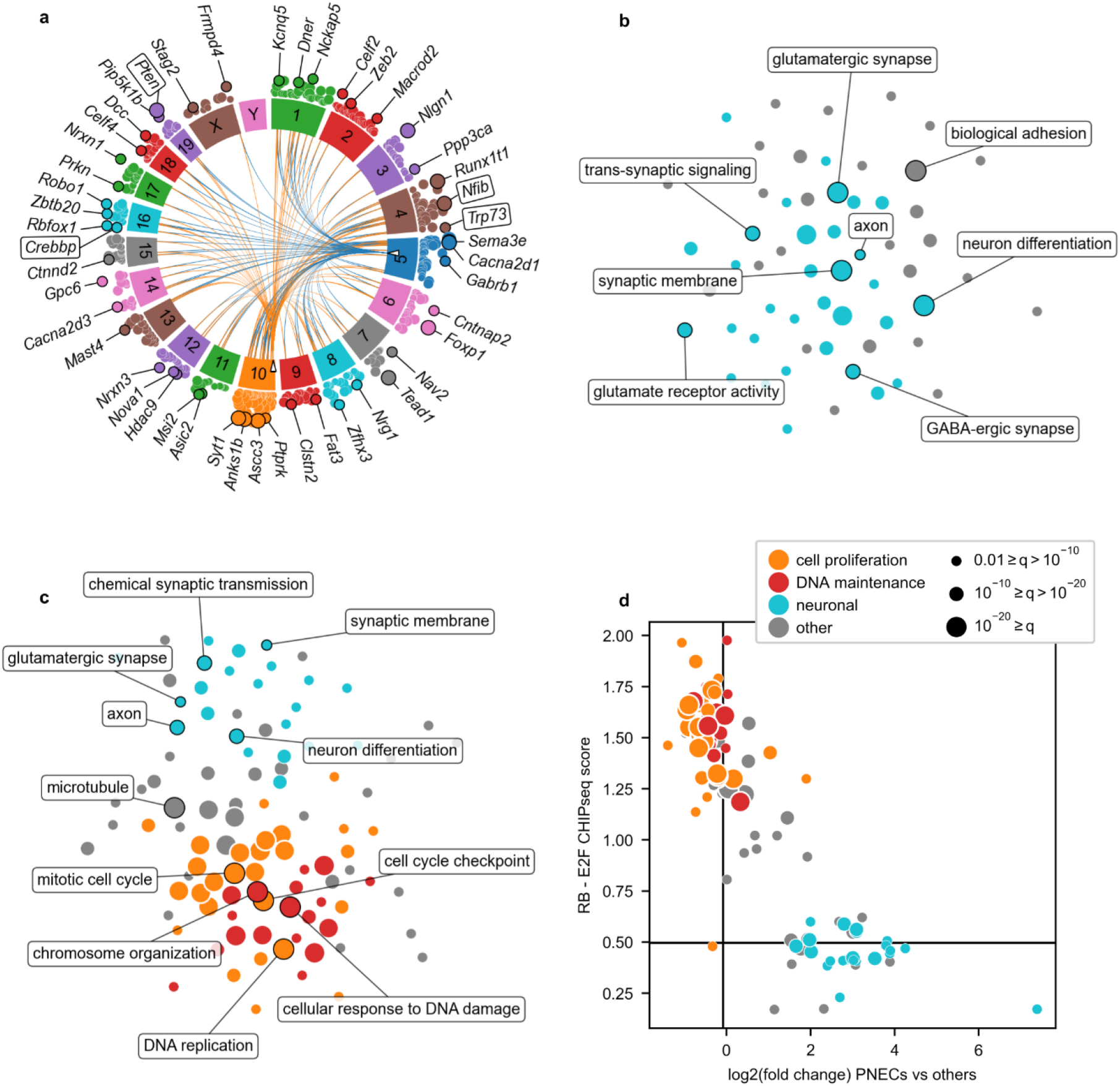
Genome-wide analysis of small cell lung cancer across species. **a)** Circos plot of the *piggyBac* screen in 303 murine tumors. The chord plot in the middle shows the transpositions from the donor loci (empty triangles) on chromosomes 5 and 10 to the 100 genes with the most significant enrichment in transposon insertions. The middle layer shows the chromosome labels. The scatter plot in the outer layer includes all genes with a significant enrichment in transposon insertions (q<0.1). Selected genes are annotated and genes previously linked to SCLC are surrounded by label boxes. **b)** Force-directed graph of gene ontology analysis, showing gene sets enriched in both the *piggyBac* dataset and the analysis of human genetic data. Most gene sets are related to synaptic and neuronal functions (light blue). **c)** Force-directed graph of gene ontology analysis, showing gene sets enriched for genes upregulated in SCLC compared to other types of cancer from the TCGA dataset and to healthy tissues from the GTEx dataset. **d)** Scatter plot of the gene sets in **c**. On the y axis is the RB-E2F score, calculated using CHIPseq data from the CISTROME database. A high score indicates strong CHIPseq signal in experiments with antibodies against RB1, RBL2, E2F1, E2F2, E2F3, E2F4, or E2F5 near the promoter of the upregulated genes included in the gene set. On the x axis is the fold change in the log2 scale of PNECs versus other lung cells in scRNA-seq data downloaded from Travaglini *et al*.^40^. A high fold change indicates that the upregulated genes within the gene set are already upregulated in healthy PNECs.

In addition to recurrent insertions in genes with an established role in SCLC biology, we also identified several genes associated with the formation of synapses. For example, *Nrxn1* (encoding neurexin-1, **Extended Data Fig. 2e**) and *Nlgn1* (encoding neuroligin-1, **Extended Data Fig. 2f**) are involved in synapse formation, where they act by binding to each other ^24–27^. *Dcc* (encoding deleted in colorectal cancer, **Extended Data Fig. 2g**), is a receptor for Netrin-1, an axon guidance cue active during neuronal development ^28^. *Reln* (encoding reelin, **Extended Data Fig. 2h**) is involved in neurodevelopment and synaptic plasticity and reelin signaling inhibits the degradation of NOTCH ^29^, which acts as tumor suppressor in SCLC ^2^. Therefore, our genetic screen recapitulates much of the known biology of SCLC and suggests a novel role for synaptic genes in SCLC.

### Comprehensive cross-species genomics analysis in murine and human SCLC confirms recurrent aberrations in synaptic gene sets

To cross-validate the recurrent transposon integration sites revealed by our *piggyBac* screen, we next assembled and re-analyzed whole exome and whole genome sequencing data from 456 specimens derived from seven distinct human SCLC datasets ^2,30–35^. This cohort is more than four times as large as the biggest individual data sets and therefore has much greater statistical power. The specimens include cell lines, primary tumors and metastases derived from both chemotherapy-naive and -exposed patients (**Extended Data Fig. 3a**). We note that these distinct SCLC samples did not show any substantial differences in their mutation profiles (**Extended Data Fig. 3b**) and contained genomic aberrations in genes with previously described roles in SCLC biology, such as *TP53, RB1, CREBBP* and *PTEN* (**Extended Data Fig. 4a-d, Extended Data Table 2**). Reassuringly, the analysis of the human data sets led to the identification of a statistically significant number of mutations in several genes that were recurrently targeted by *piggyBac* transposon integrations, including *NRXN1, NLGN1, DCC* and *RELN* (**Extended Data Fig. 4e-h**). Overall, the *piggyBac* and human data sets were highly overlapping (p = 7.2*10^-37^, Fisher’s exact test).

To functionally categorize the composition of the genes detected in the human sequencing data sets and in the *piggyBac* screen, we first asked which gene ontology (GO) gene sets ^36,37^ are significantly enriched in both data sets (**Fig. 1b, Extended Data Tables 3, 4**). Unexpectedly, the vast majority of the enriched terms were related to neuronal phenotypes and synaptic functions, such as *Synaptic Membrane, Glutamatergic Synapse, Glutamate Receptor Activity* and *Transsynaptic Signaling*, among others. Therefore, the only clear genetic signal we identified at the network level in 456 human tumors and 303 murine tumors is related to neuronal and synaptic functions.

### Specific expression of synaptic and neuronal gene sets is conserved between SCLC cells and PNECs

To further probe the relevance of these recurrent aberrations and *piggyBac* insertions in synaptic genes, we next analyzed transcriptome data derived from tumor specimens and normal tissue. For that purpose, we collected raw expression data from the George *et al*.^2^ and Rudin *et al*.^35^ data sets and reanalyzed them using the TCGA transcriptome pipeline, in order to identify gene sets with expression levels that are enriched in SCLC transcriptomes, compared to 33 distinct cancer entities, including lung adenocarcinoma, lung squamous cell carcinoma, pheochromocytoma, pancreatic ductal adenocarcinoma, diffuse large B cell lymphoma and others (**Fig. 1c, Extended Data Fig. 5, Extended Data Tables 5, 6**). We similarly deployed the GTEx pipeline, to ask which gene sets are specifically enriched in SCLC transcriptomes, compared to those derived from 27 healthy tissues (**Fig. 1c, Extended Data Fig. 5, Extended Data Tables 7, 8**). Using this approach, we identified several gene sets involved in DNA replication, cell cycle checkpoint signaling, chromosome organization and the DNA damage response (**Extended Data Fig. 5a-d**). These gene sets included individual genes such as *CCNE2*, encoding cyclin E2, which is a central driver of the G_1_S phase transition induced upon RB1 repression and *TOP2A*, encoding topoisomerase 2, the drug target of etoposide, which constitutes a central pillar of our current SCLC frontline regimens ^1,38^. Individual genes identified in the *piggyBac* and human genetic datasets were highly expressed, such as *NRXN1, NLGN1, DCC* and *RELN* (**Extended Data Fig. 5e-h**). Importantly, we also identified several of the very same gene sets that were enriched at the genetic level, such as *Synaptic Membrane, Glutamatergic Synapse, Chemical Synaptic Transmission* and *Neuron Differentiation*, among others (**Fig. 1c**). To further characterize the gene sets that are specifically enriched in transcriptomes of SCLC tumors, we derived an RB-EF2 score, using ChIP-Seq data from the CISTROME database ^39^. A high score indicates a strong, ChIP-Seq-verified presence of RB1, RBL2, E2F1, E2F2, E2F3, E2F4 or E2F5 near the promoter of the upregulated genes included in the gene set. As a cross-reference, we plotted gene expression profiles derived from PNECs versus other lung-resident cell populations on a log2 scale, deploying a previously published data set ^40^, with a high fold change indicating that the upregulated genes within a given gene set are specifically upregulated in PNECs. This analysis indicated that the SCLC-specific expression of neuronal and synaptic gene sets is conserved from the PNEC-derived SCLC cell of origin, whereas the high expression levels of genes associated with cell cycle regulation and genome maintenance appear to be largely driven by RB-E2F signaling (**Fig. 1d**). Therefore, two signals are evident in human SCLC at the expression level. First, the high expression of cell cycle gene sets downstream of the RB-E2F axis. Second, the high expression of neuronal and synaptic gene sets inherited from the cell of origin, which are strikingly overlapping with the GO terms we identified at the genetic level.

### SCLC cells receive functional synaptic inputs

The observation that gene sets related to neuronal phenotypes and synapses constitute the strongest and most consistent signal in our *piggyBac* screen and in human samples prompted us to further investigate a potential physical interaction between SCLC tumor cells and neurons. As a first step, we established co-cultures *in vitro* and examined the behavior of murine cortical neurons following addition of the human SCLC cell lines COR-L88 and DMS273. Within five days of co-culturing, we consistently observed clusters of SCLC cells that were profusely contacted by neuronal axons bearing immunoreactivity for the vesicular glutamate transporter 1 (VGLUT1), a pre-synaptic glutamatergic marker (**Fig. 2a**). Analysis of these putative axon-SCLC cell contact regions confirmed the presence of numerous VGLUT1-positive boutons decorating the border of the cancer cells (**Fig. 2a**).

**Figure 2:**
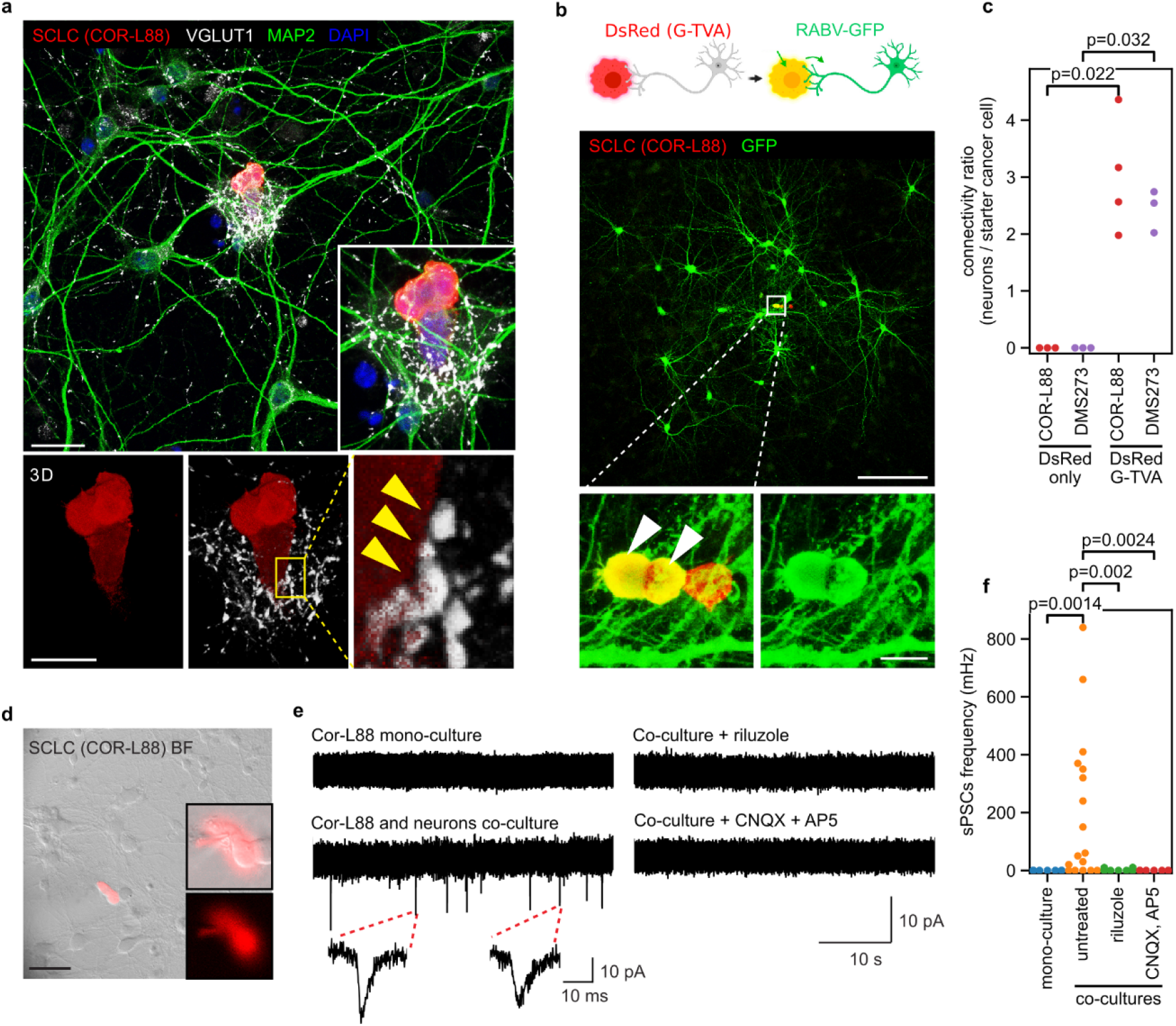
SCLC cells form functional synapses. **a)** Co-culture of neurons (immunolabeled against MAP2) and SCLC cells (COR-L88, expressing DsRed) showing the appearance of dense VGLUT1-positive punctae onto SCLC cells contacted by neuronal terminals. Bottom panels show a 3D reconstruction of the inset in the main panel. Bars, 20 and 15 μm. **b)** Strategy utilized to retrogradely trace neurons monosynaptically connected with SCLC cells engineered to express DsRed, G and TVA. Addition of EnvA-pseudotyped RABV-GFP resulted in primary infection of SCLC cells and the secondary emergence of nearby GFP-only-positive neurons. Bottom panels show enlarged views of the boxed area containing double-positive starter SCLC cells. Bars, 100 and 10 μm. **c)** Connectivity ratio per starter cell for 2 SCLC cell lines in the presence or absence of G and TVA expression (n = 3-4 experiments). **d)** Example of a patched DsRed-expressing SCLC cell under whole-cell configuration in neuron-SCLC co-cultures. Bar, 30 μm. **e)** Representative whole-cell, voltage-clamp recordings of sPSCs in SCLC cells in the presence or absence of neurons, with or without addition of riluzole or synaptic blockers. **f)** Quantification of sPSC frequency under the conditions illustrated in **e** (n = 7-17 cells per condition).

To assess the functionality of these putative synaptic contacts between neurons and cancer cells, we next conducted retrograde monosynaptic tracing experiments. For this purpose, we employed a replication-incompetent EnvA-pseudotyped RABV-GFP, which can only infect cells expressing the avian viral receptor TVA. Additionally, only cells that express the glycoprotein (G) can transmit the virus to first-order pre-synaptic cells ^41^. We transduced SCLC cells with a DsRed retrovirus encoding for the avian viral receptor TVA and the glycoprotein (G) and co-cultured them with cortical neurons (**Fig. 2b**). Upon establishment of SCLC-neuron coculture, RABV-GFP was added to the media. Intriguingly, we often found RABV-GFP SCLC cells being surrounded by clusters of GFP-positive neurons (**Fig. 2b, Extended Data Fig. 6a,b**), indicative of retrograde RABV spread from SCLC “starter” cells into synaptically connected neurons. We also found strong VGLUT1 immunoreactivity in RABV-infected neuronal fibers impinging onto infected cancer cells in these samples (**Extended Data Fig. 6c**). Analysis of cultures containing COR-L88 or DMS273 cells revealed an average connectivity ratio of more than two neurons per “starter” cancer cell. In contrast, no neurons were labeled when the SCLC cells were engineered to express DsRed but not the viral receptor TVA or the glycoprotein G (**Fig. 2c**).

To further substantiate these findings, we next assessed by electrophysiology whether SCLC cells can establish functional synapses with neurons. While wholecell patch-clamp recordings of COR-L88 mono-cultures revealed an absence of spontaneous inputs, the same cells developed spontaneous post-synaptic currents (sPSCs) when co-cultured with cortical neurons (**Fig. 2d-e**). Importantly, these currents could be blocked with either a cocktail of AMPA and NMDA receptor antagonists (CNQX and AP5) or with the glutamate release inhibitor 2-amino-6-trifluoromethoxy benzothiazole (riluzole) ^42^ (**Fig. 2e-f**).

We next investigated whether these findings could be validated *in vivo* by conducting immunohistochemical analyses of lung sections isolated from tumor-bearing *Rb1^fl/fl^;Trp53^fl/fl^* mice in combination with calcitonin gene-related peptide (CGRP) immunofluorescence, which labels normal PNECs, SCLC tumors and sensory fibers derived from the dorsal root, jugular and nodose ganglia ^16^ (**Extended Data Fig. 7**). Interestingly, SYP-, PGP9.5-, GAP43-, VGLUT1-, and P2X3-positive nerve fibers were also detectable in a subset of healthy PNECs, clustered into neuroepithelial bodies (NEBs), and in small SCLC tumors (**Fig. 3a-c**; **Extended Data Fig. 8a-h**). In contrast, larger tumors were mostly devoid of intra-lesional nerve fibers and, when present, GAP43-positive fibers were observed at the tumor border (**Fig. 3c; Extended Data Fig. 8h**). The presence of nerve fibers within small *Rb1^fl/fl^; Trp53^fl/fl^*-derived SCLC tumors was further corroborated at the ultrastructural level with electron microscopy, where vesicle-enriched axon-like fibers were found in contact with tumor cells (**Extended Data Fig. 8i-l**). Biopsies from human SCLC patients in advanced stage showed a pattern comparable to large murine tumors, with nerve fibers immunoreactive for neurofilaments and synaptophysin located at the border or in the vicinity of the tumors (**Extended Data Fig. 9**).

**Figure 3:**
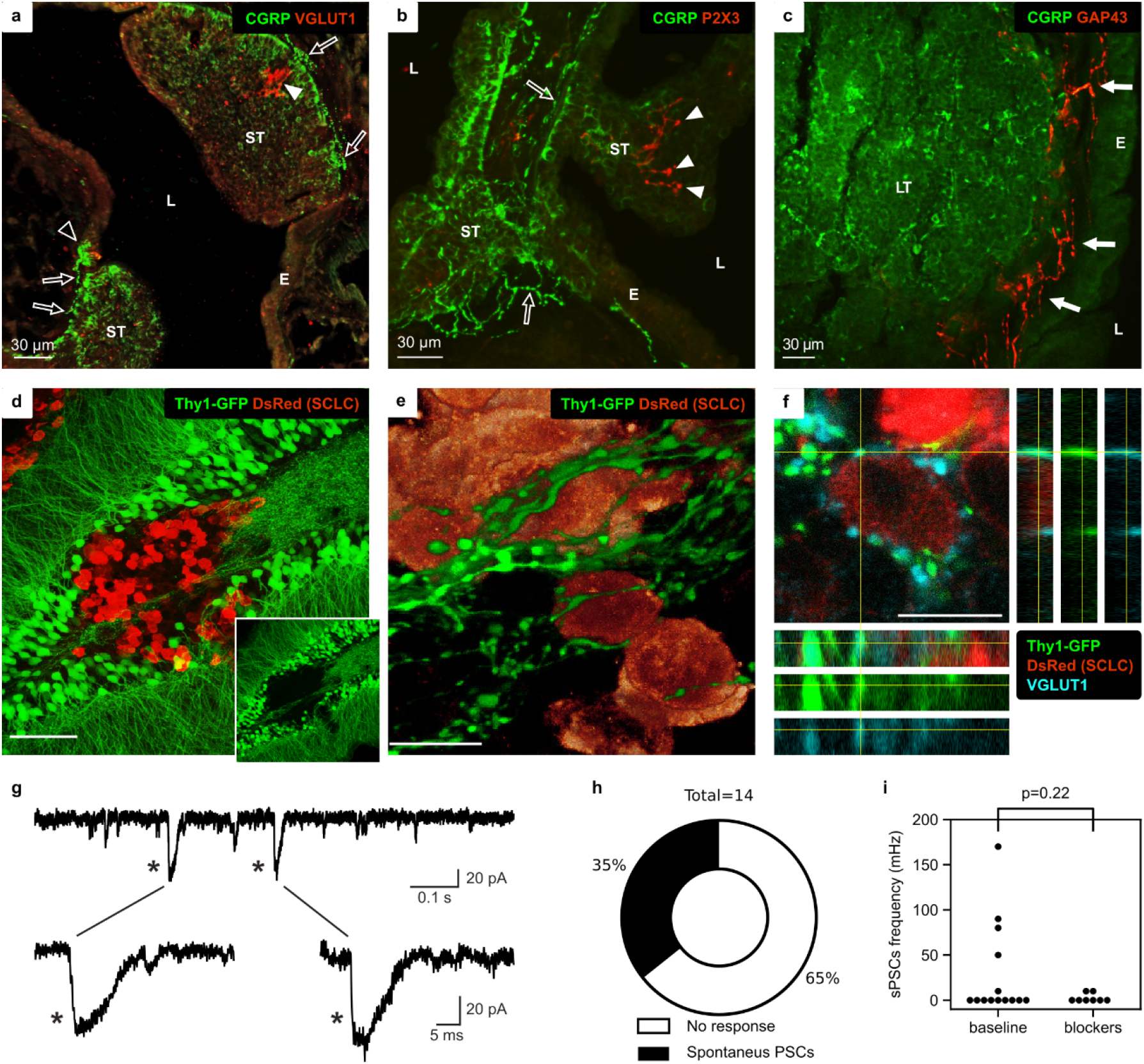
Innervation of murine SCLC tumors. **a)** Confocal image of an intrapulmonary airway of a *Rb1^fl/fl^;Trp53^fl/fl^* mouse. Two small tumors (ST) and a normal NEB (open arrowhead) are visualized with CGRP (FITC, green) and can be observed to bulge in the airway lumen (L). VGLUT1-ir nerve terminals are detected contacting the NEB and arborising (arrowhead) within one of the tumors (Cy3, red). CGRP-ir nerve fibers (open arrows) can be observed at the base of the tumors and NEB. E: epithelium. **b)** P2X3-positive nerve terminals (arrowheads, red Cy3) can be seen to arborise between the CGRP-ir neuroendocrine cells of a small tumor (ST). CGRP-ir nerve fibers (open arrows) can be observed at the base of the tumor. L: lumen; E: epithelium. **c)** Immunolabelling of a CGRP-ir large tumor (LT, green FITC). The connective tissue between the tumor and the epithelium (E) harbors many GAP43-ir nerve fibers (arrows, red Cy3), which do not appear to penetrate the tumor mass. L: lumen. **d)** Immunolabeling of SCLC cells (DsRed) transplanted into the hippocampus of Thy1-GFP mice. Insets shows the core tumor area being devoid of GFP-positive fibers. Bar, 70 μm. **e)**3D reconstruction of SCLC cells located in the tumor periphery being surrounded by GFP-positive axonal boutons. Bar, 10 μm. **f)** Co-localization analysis of GFP- and VGLUT1-positive boutons contacting a DsRed-expressing SCLC cell. Bar, 10 μm. **g)** Representative whole-cell, voltage-clamp recordings of grafted DsRed-expressing SCLC cells in acute brain slices. **h-i)** Quantification of cancer cells displaying sPSCs sensitive to synaptic receptor blockers (CNQX, AP5 and Bicuculline).

To experimentally test for the formation of functional synaptic contacts between SCLC cells and neurons *in vivo*, we next performed allograft experiments into the brain of healthy *Thy1-GFP* transgenic mice. For that purpose, we transplanted DsRed-expressing RP cells into the hippocampus of recipient mice, in which excitatory neuronal subsets express EGFP. Using confocal microscopy analysis of brain sections, we found that by 10-12 days after transplantation the cancer cells located in the periphery of the tumor were profusely contacted by GFP-positive boutons and axonal bundles (**Fig. 3d-f**). Moreover, most of these GFP-positive boutons were strongly immunoreactive for VGLUT1 (**Fig. 3f**). In stark contrast, GFP-positive nerve fibers appeared to be largely depleted from the central regions of the tumor mass (**Fig. 3d**), consistent with our observations in large autochthonous murine tumors and patient biopsies and suggesting a putative neurotoxic effect within this area. Whole-cell patch-clamp recordings of transplanted SCLC cells in acute brain slice preparations revealed detectable biphasic sPSCs in 35% of cells (n = 14 cells from 6 mice) (**Fig. 3g,h**). Of note, pre-treatment of the brain slices with a mixture of synaptic blockers (CNQX, AP5 and Bicuculline) was sufficient to abolish most PSCs (**Fig. 3i**), suggesting that transplanted SCLC cells engage in synaptic transmission in brain tissue. Together, these experiments indicate that SCLC cells are capable of forming *bona fide* synapses with neurons *in vitro* and *in vivo*.

### Cortical neurons enhance the proliferation kinetics of human SCLC cells

To understand the implications of synaptic contacts for SCLC cell behavior, we next tested whether SCLC cells derive a growth advantage when kept in co-culture with neurons, as compared to mono-cultures. To this end, we compared the proliferative capacity of DsRed-expressing human COR-L88 cells when seeded at low density and maintained either alone (mono-culture) or in co-culture with cortical neurons, followed by EdU labeling 2h prior to analysis. While only few scattered EdU-positive SCLC cells were found in mono-cultures, SCLC cells in co-cultures often appeared as significantly larger proliferating clusters (**Fig. 4a-d**). Consistent with these experiments, the presence of neurons caused a significant increase in the proliferation rate of five distinct human SCLC cell lines (COR-L88, DMS273, H146, H69, H524), compared to the mono-cultures (**Fig. 4e**). These data demonstrate that SCLC cells derive a growth benefit when proliferating in close proximity to neurons.

**Figure 4:**
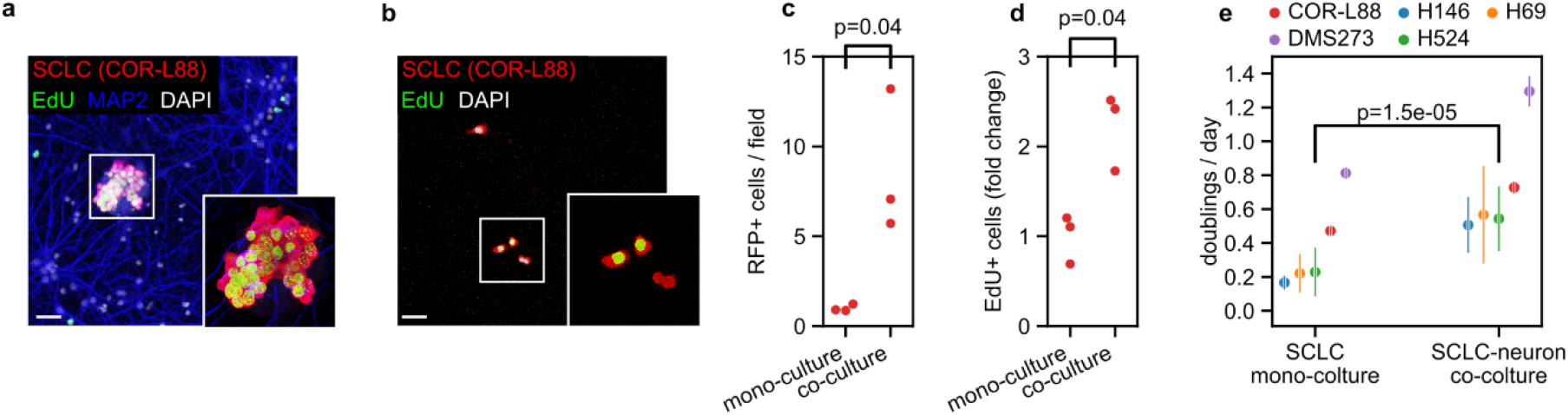
Enhanced proliferation of SCLC cells in the presence of neurons. **a)** Example of SCLC cells (COR-L88, DsRed) cultured in presence of neurons and following treatment with EdU to label dividing cells. Bar, 30 μm. **b)** Example of COR-L88 cells cultured in the absence of neurons and following treatment with EdU to label dividing cells. Bar, 30 μm **c)** Quantification of CORL-88 cells in the absence or presence of neurons (n = 3 experiments, Mann–Whitney test). **d)** Quantification of EdU-positive CORL-88 cells in absence or presence of neurons (n = 3 experiments, Mann–Whitney test). **e)** Proliferation speed of SCLC cell lines in mono-culture or in co-culture with mouse cortical neurons. Doublings per day were calculated between day 3 and 6 after seeding. The q value is the Wilcoxon rank sum p value adjusted for false discovery rate. Each point is the median of three to four independent experiments each performed in triplicate. The vertical lines are the median absolute deviation from the median.

### Pharmacological perturbation of the glutamatergic system displays therapeutic efficacy in an autochthonous SCLC mouse model

Given the identification of the GO term *Glutamatergic Synapse* in all of our data sets (**Fig. 1**), the formation of functional synaptic contacts between SCLC cells and glutamatergic murine neurons *in vitro* and *in vivo* (**Fig. 2a, Fig. 3f**) and the presence of glutamatergic nerve fibers within a subset of primary SCLC tumors (**Fig. 3a**), we next sought to target the glutamatergic system therapeutically. The SCLC samples that we analyzed at the expression level can be broadly classified into classic SCLC with strong neuroendocrine features and variant SCLC with lower expression of neuroendocrine features, based on the expression levels of the neuroendocrine marker genes *SYP*, *CHGA, ENO2, NCAM1, S100B* (**Fig.5a**). The expression of genes within the GO term *Glutamatergic Synapse* was particularly high in classic SCLC samples in general and in the *ASCL1*- and *NEUROD1-* expressing subtypes, in particular ^4^ (**Fig. 5b**), suggesting that these subtypes might particularly benefit from the manipulation of the glutamatergic system.

**Figure 5:**
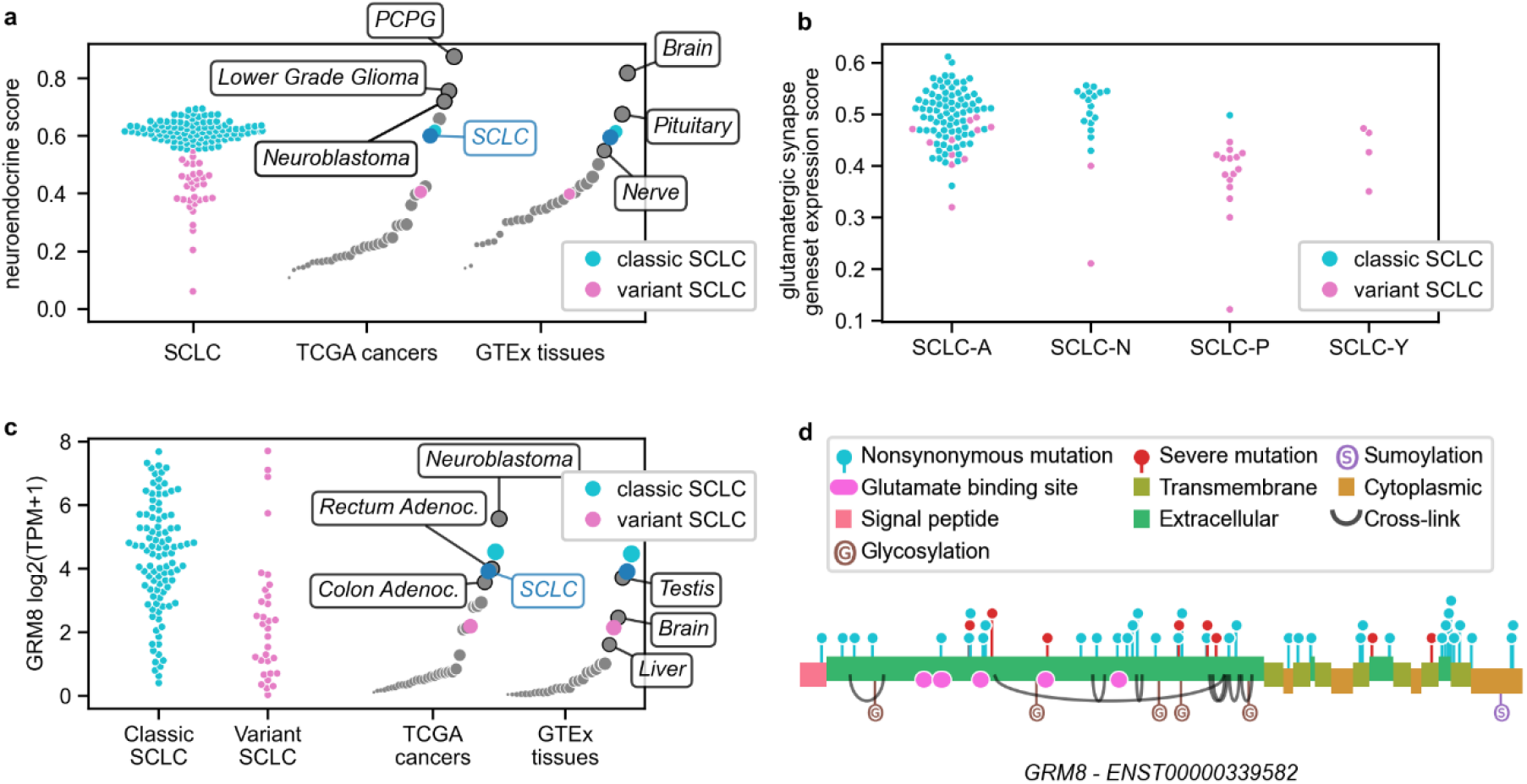
*GRM8* is a potential target in SCLC subtypes with strong neuroendocrine features. **a)** Individual SCLC samples can be separated into two populations based on the expression of neuroendocrine markers (*SYP, CHGA, ENO2, NCAM1, S100B*) assembled into a neuroendocrine score. Compared to other cancers and healthy tissues, SCLC shows a high neuroendocrine score. PCPG: pheocromocytoma and paraganglioma **b)** SCLC samples are divided into subgroups based on the expression of four transcription factors: *ASCL1* (SCLC-A), *NEUROD1* (SCLC-N), *POU2F3* (SCLC-P) and *YAP1* (SCLC-Y). Samples with a classic, highly neuroendocrine phenotype correspond to SCLC-A and SCLC-N subtypes and express higher level of genes included in the GO term *Glutamatergic Synapse* **c)** The expression of *GRM8* is higher in SCLC than in most other cancers and tissues and is especially high in classic SCLC with strong neuroendocrine features. **d)** The GRM8 protein is shown, with annotations from the UniProt Knowledgebase. Mutations identified in SCLC samples are shown as a lollipop chart above the protein. Severe mutations (stop, frameshift, start-loss, and canonical splice-site mutations) are shown in red. Nonsynonymous mutations (amino-acid substitution, non-frameshift indels, non-canonical splice site mutations) are shown in light blue.

Among the possible molecular targets within this system are the glutamate receptors, which we also identified as individual genes targeted by transposon insertions in our *piggyBac* screen (*GRID1, GRIK2, GRIN3A, GRM1, GRM3, GRM5, GRM8*), in human mutation data (*GRIA1, GRIA2, GRIA3, GRIA4, GRID2, GRIK2, GRIK3, GRIK4, GRIN2A, GRIN2B, GRIN3A, GRM1, GRM3, GRM5, GRM8*) and at the expression level (*GRIA2*, *GRIN3A*, *GRIK3, GRIK5, GRM2, GRM4, GRM8*). Prominent among them was *GRM8*, an inhibitory metabotropic glutamate receptor that has been shown to counteract glutamate signaling by negatively regulating cAMP-dependent sensitization of inositol 1,4,5-trisphosphate receptors, thereby limiting glutamate-induced calcium release from the ER ^43^. In our datasets, *GRM8* shows specific expression in SCLC and a few other tumors types (**Fig. 5C**) and a statistically significant enrichment of both non-synonymous mutations and more severe loss of function mutations (stop, frameshift, start-loss, and canonical splicesite mutations) (**Fig. 5D**). Re-analysis of scRNA-seq data from Chan *et al*. ^44^ confirmed that GRM8 is specifically expressed in the cancer cells of SCLC-A and SCLC-N subtypes (**Extended Data Fig. 10**). The specific expression suggests that GRM8 can be targeted, while the enrichment in loss of function mutations suggests that its activity is detrimental to the tumor. Based on these data, we selected (S)-3,4-Dicarboxyphenylglycine (DCPG) and riluzole, two compounds with predicted anti-glutamatergic effects, for preclinical testing. DCPG is a potent and selective agonist of GRM8, with a reported IC_50_ of 31 nM and 100-fold specificity for GRM8 over other glutamate receptors ^45^. Riluzole is an FDA-approved inhibitor of glutamate release, which could inhibit the spontaneous post-synaptic currents in our co-culture experiments (**Fig. 2e,f**).

To evaluate the efficacy of DCPG and riluzole *in vivo*, we induced SCLC tumors in RP mice via intratracheal Ad-CMV-Cre instillation. Upon MRI imaging-verified tumor formation, animals were exposed to DCPG or riluzole as single agents or their relative vehicles. The response was evaluated by longitudinal MRI imaging every two weeks. While all tumors from vehicle-treated mice progressed during treatment, the response was significantly improved both in the DCPG cohort (q=7.7*10^-5^) and in the riluzole cohort (q=7.7*10^-5^), with several tumors showing short term stable disease or slower growth and a small subset of tumors showing modest shrinkage (**Fig. 6a**). The DCPG- and riluzole-treated mice also showed a significantly improved survival, with a median survival of 66 days (DCPG) and 71.5 days (riluzole), compared to 54 days in the controls (**Fig. 6b**). We then compared animals treated with frontline cisplatin/etoposide chemotherapy with mice that received chemotherapy with the addition of DCPG or riluzole. Tumors exposed to chemotherapy showed a mixed response, which included shrinkage, stable disease and progressive disease. In the context of chemotherapy, the inclusion of DCPG did not significantly improve the response of SCLC tumors (**Fig. 6c**). In contrast, the combination of chemotherapy and riluzole resulted in a significantly improved response, with almost all tumors displaying partial response or stable disease (q=0.0086, **Fig. 6c**). Similarly, the inclusion of DCPG into the frontline chemotherapy regimen in our mice did not result in significantly improved survival (**Fig. 6d**), while the inclusion of riluzole resulted in significant and substantial improvement of 21 days (p = 0.0097).

**Figure 6:**
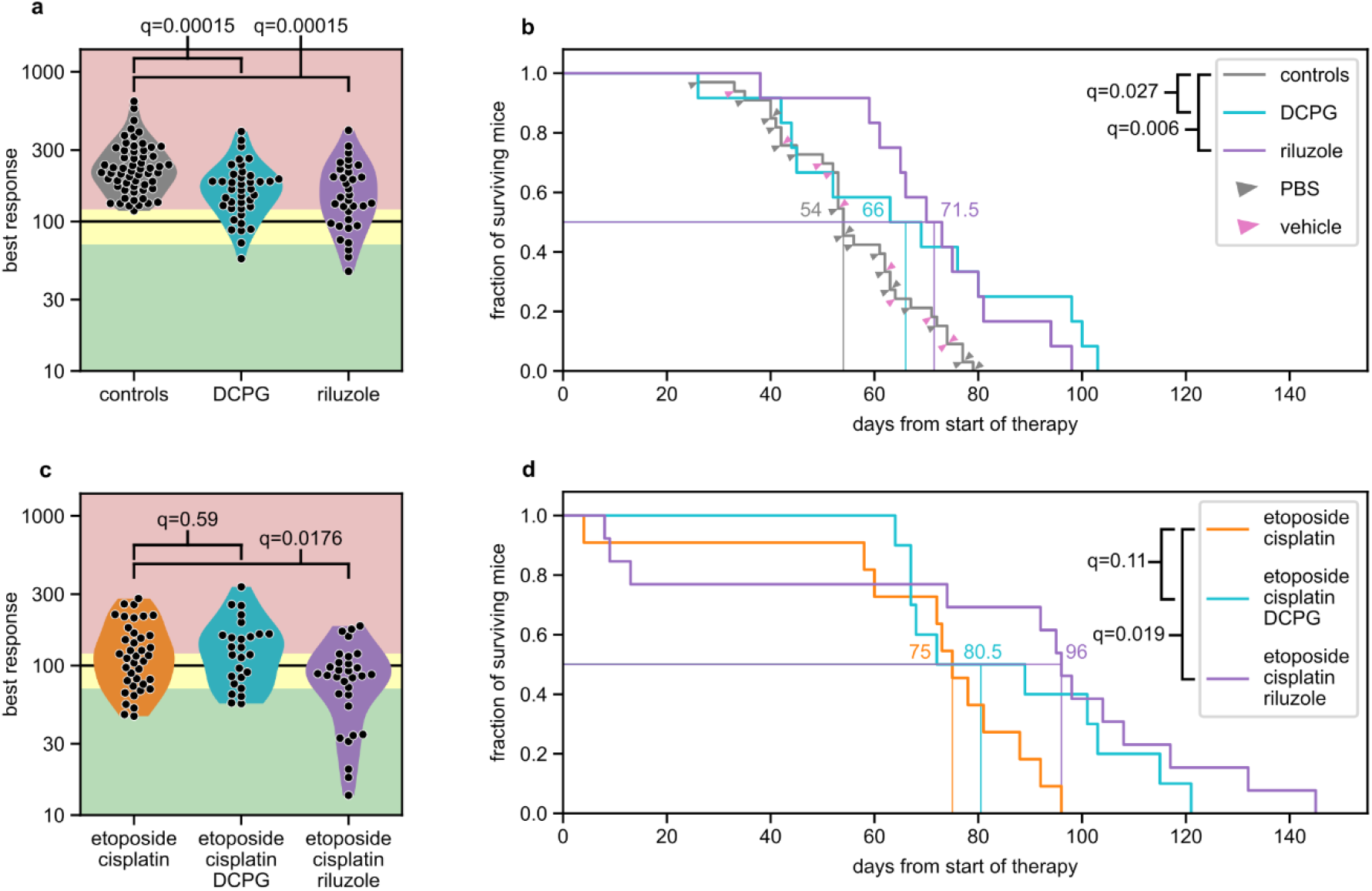
Anti-glutamatergic drugs are effective in small cell lung cancer. **a)** Best response of individual tumors from *Rb1^fl/fl^;Trp53^fl/fl^* mice treated with DCPG, riluzole or with their relative controls (PBS or vehicle solution). The best response is shown as percentage relative to the tumor size in the last MRI scan before therapy. The red area indicates progression (>120%), the yellow area indicates stable disease (70-120%) and the green area indicates response (<70%). Treatment with DCPG or riluzole results in a significantly improved response, mainly characterized by slower growth or stable disease. **b)** survival of the mice in **a**. Treatment with DCPG (n = 12 mice) or riluzole (n = 12 mice) results in significantly longer survival compared to control mice (n = 23 for PBS plus n = 10 for riluzole vehicle). Log-rank test with FDR correction. **c)** best response of individual tumors from *Rb1^fl/fl^;Trp53^fl/fl^* mice treated with chemotherapy (etoposide and cisplatin every two weeks) with and without riluzole or DCPG. Adding riluzole to chemotherapy results in a significantly improved response, with almost all tumors showing either response or stable disease. In contrast, adding DCPG to chemotherapy does not result in improved response **f)** survival of the mice in **e**. The combination of riluzole and chemotherapy (n = 13 mice) results in significantly longer survival than chemotherapy alone (n = 11 mice), while the improvement in survival caused by the addition of DCPG to chemotherapy (n = 10 mice) is not statistically significant. Log-rank test with FDR correction.

Therefore, anti-glutamatergic compounds display pre-clinical anti-SCLC activity alone and in combination with an established frontline chemotherapy regimen *in vivo*.

## Discussion

Here, we performed a large *in vivo* genetic screen using the *piggyBac* transposition system in a mouse model of SCLC and cross-validated our findings with the reanalyses of the available genetic and expression data from human SCLC. Surprisingly, almost all gene sets we identified at the genetic level are related to a neuron-like phenotype in general and to synapses and glutamatergic signaling in particular.

Our co-culture experiments revealed a striking ability of SCLC cells to receive functional glutamatergic inputs and transfer RABV to pre-synaptic neurons, demonstrating the formation of synaptic contacts. We further confirmed these findings *in vivo*, using transplantation of SCLC cells into the brain of recipient mice. These data reveal that *bona fide* synapses can form between neurons and cancer cells that do not derive from the nervous system.

Similar to the overall expression of synaptic genes, we speculate that the ability to form synapses is inherited from the PNECs. Consistent with this notion, we could detect fibers known to innervate PNECs, such as VGLUT1-positive vagal fibers and P2X3-positive fibers from the dorsal root ganglia ^16^, within a subset of small SCLC tumors in the autochthonous RP mouse model of SCLC.

In line with a putative oncogenic role of SCLC-neurons interactions, all SCLC cell lines we tested derived a growth advantage when cultured in the presence of neurons. While we observed direct synaptic communication in our co-cultures, we cannot exclude a paracrine component to this proliferative advantage. In fact, we observed several CGRP-positive and GAP43-positive nerve fibers around autochthonous RP tumors, which could potentially engage in paracrine communication with the tumors. A systematic exploration and cataloguing of synaptic and non-synaptic (peri-synaptic and paracrine neuro-ligand signaling) interactions between neurons and SCLC cells will constitute an important next step forward.

Expression of genes included in the GO-term *Glutamatergic Synapse*, as well as *GRM8* expression was particularly high in human SCLC cases of the SCLC-A and SCLC-N subtypes, indicating that particularly those tumors with preserved neuroendocrine differentiation might be amenable to therapeutic interception of glutamate signaling. As SCLC is a cancer entity characterized by a high degree of inter- and intratumor heterogeneity and plasticity ^4,46–50^, the general exploitability of glutamate targeting strategies and the potential therapy sequencing algorithms remain to be defined. Taken together, our data suggest that SCLC is capable of hijacking neuronal programs, such as the ability to form functional synapses, to derive a growth advantage. As we show for anti-glutamatergic drugs, the investigation of these neuronal phenotypes may hold the key to finally providing more effective therapeutic options to SCLC patients.

## Extended Data Figures

**Extended Data Figure 1:**
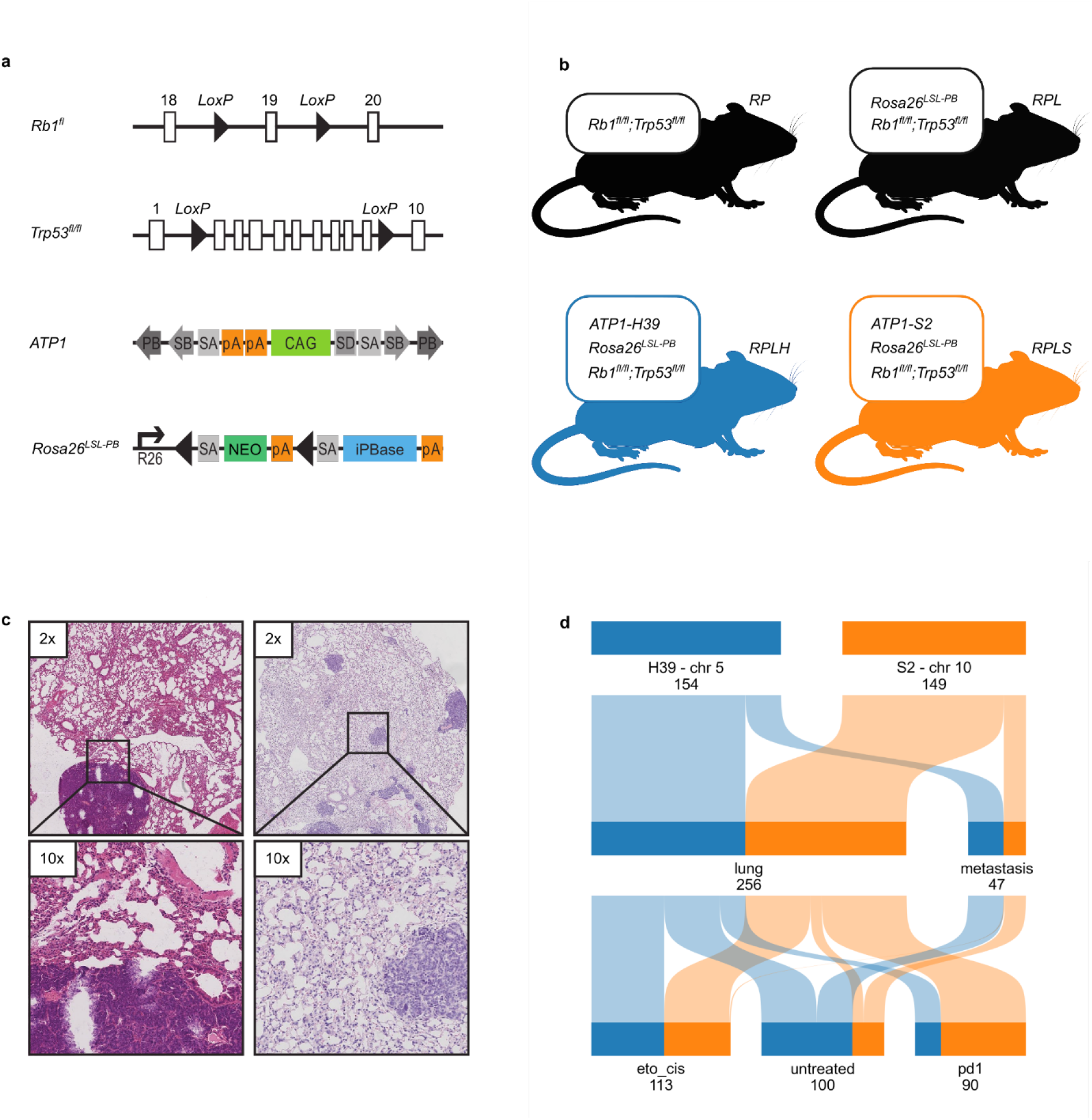
Mice used for the transpositional mutagenesis screen with the *piggyBac* system. **a)** Alleles included in the mouse model **b)** Mouse lines included in the screen carry the *Rb1^fl/fl^* and *Tp53^fl/fl^* alleles with the addition of the conditional allele to express the *piggyBac* transposase (*Rosa26^LSL-PB^*). The RPLH line (blue) additionally carries the donor allele *ATP1-H39*, with 80 copies of the *ATP1* transposon on chromosome 5. The RPLS line (orange) additionally carries the donor allele *ATP1-S2*, with 20 copies of the *ATP1* transposon on chromosome 10. **c)** Tumors derived from RPLH and RPLS mice display typical SCLC morphology.**d)** Tumors harvested from RPLH (blue) and RPLS (orange) mice include lung and metastatic samples and derive from untreated mice, from mice treated with etoposide and cisplatin and from mice treated with anti-PD1 antibody RPM1-14.

**Extended Data Figure 2:**
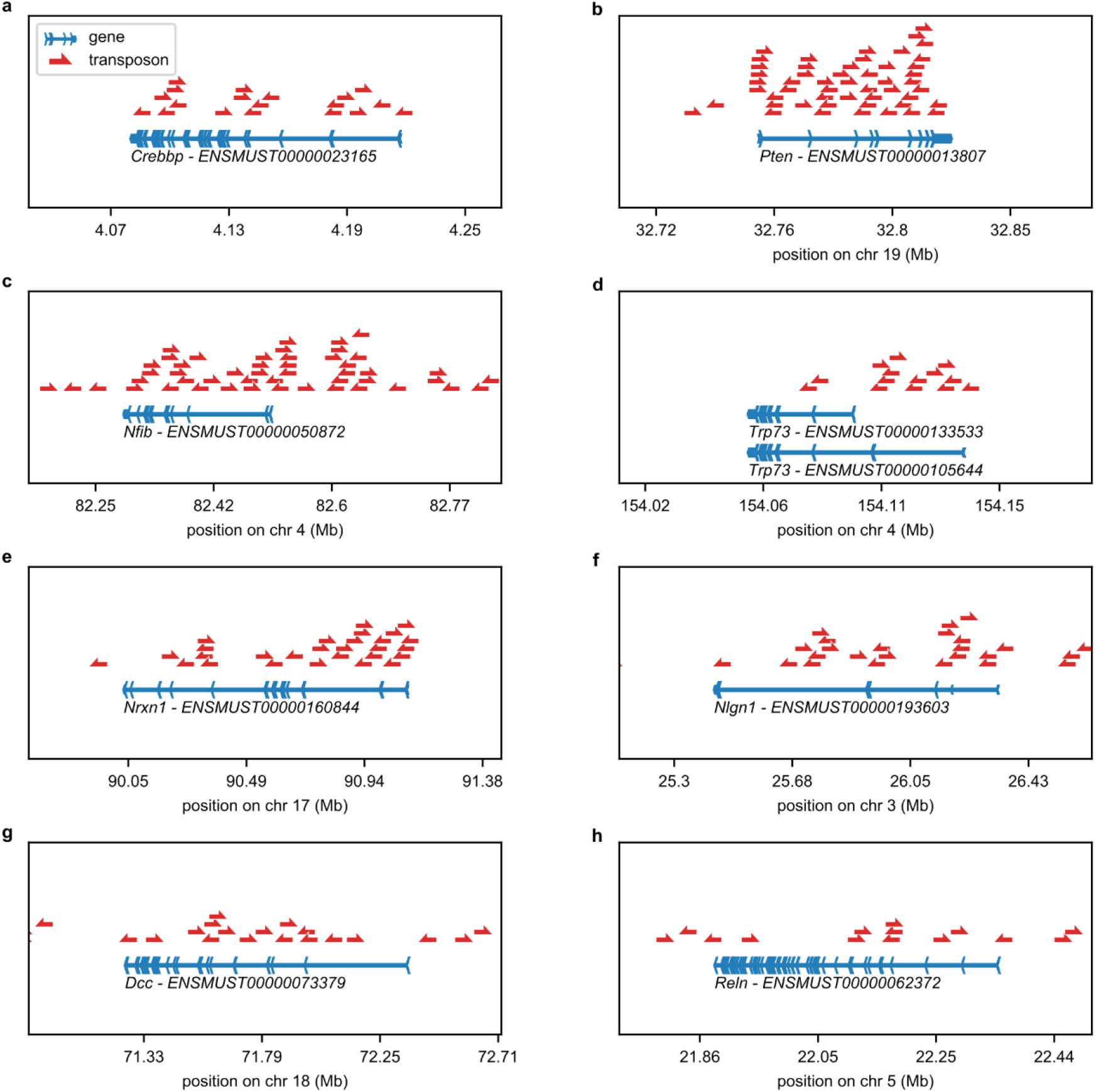
Selected genes with high rate of PiggyBac insertions. Transposon insertions (red arrows) identified in selected genes (horizontal blue lines). The orientation of the exons (vertical obtuse blue angles) point to the direction of transcription. **a)** Insertions in *Crebbp* are located within the gene and are oriented independently of the direction of transcription in a pattern that suggests a tumor suppressor function. **b)** Insertions in *Pten* are located within the gene and are oriented independently of the direction of transcription in a pattern that suggests a tumor suppressor function. **c)** Insertions in *Nfib* are located both within the gene and upstream of the gene and are oriented independently of the direction of transcription. **d)** Insertions in *Trp73* are located mainly on the long isoform and for the most part point in the direction of transcription, suggesting a tumor suppressive function for the long isoform and an oncogenic function for shorter isoforms. **e)** Insertions in *Nrxn1* are located within the gene and are oriented independently of the direction of transcription in a pattern that suggests a tumor suppressor function. **f)** Insertions in *Nlgn1* are located within the gene and are oriented independently of the direction of transcription in a pattern that suggests a tumor suppressor function. **g)** Insertions in *Dcc* are located within the gene and are oriented independently of the direction of transcription in a pattern that suggests a tumor suppressor function. **h)** Insertions in *Reln* are located within the gene and are oriented independently of the direction of transcription in a pattern that suggests a tumor suppressor function.

**Extended Data Figure 3:**
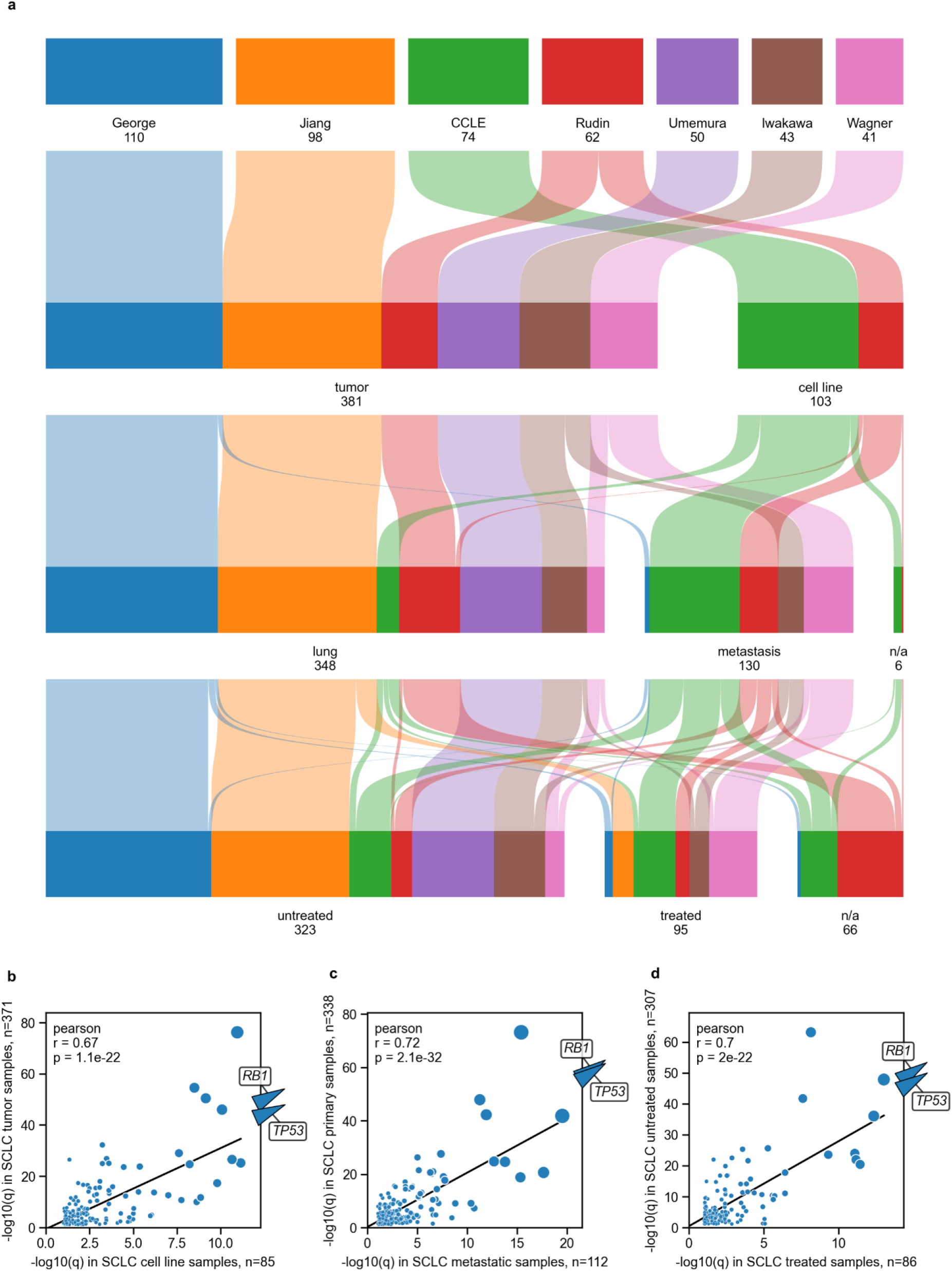
Overview of genetic data from human patients. **a)** Origin and characteristics of human samples from different studies. **b)** Similar genes are identified in tumor and cell line samples. **c)** Similar genes are identified in primary and metastatic samples. **d)** Similar genes are identified in treated and untreated samples

**Extended Data Figure 4:**
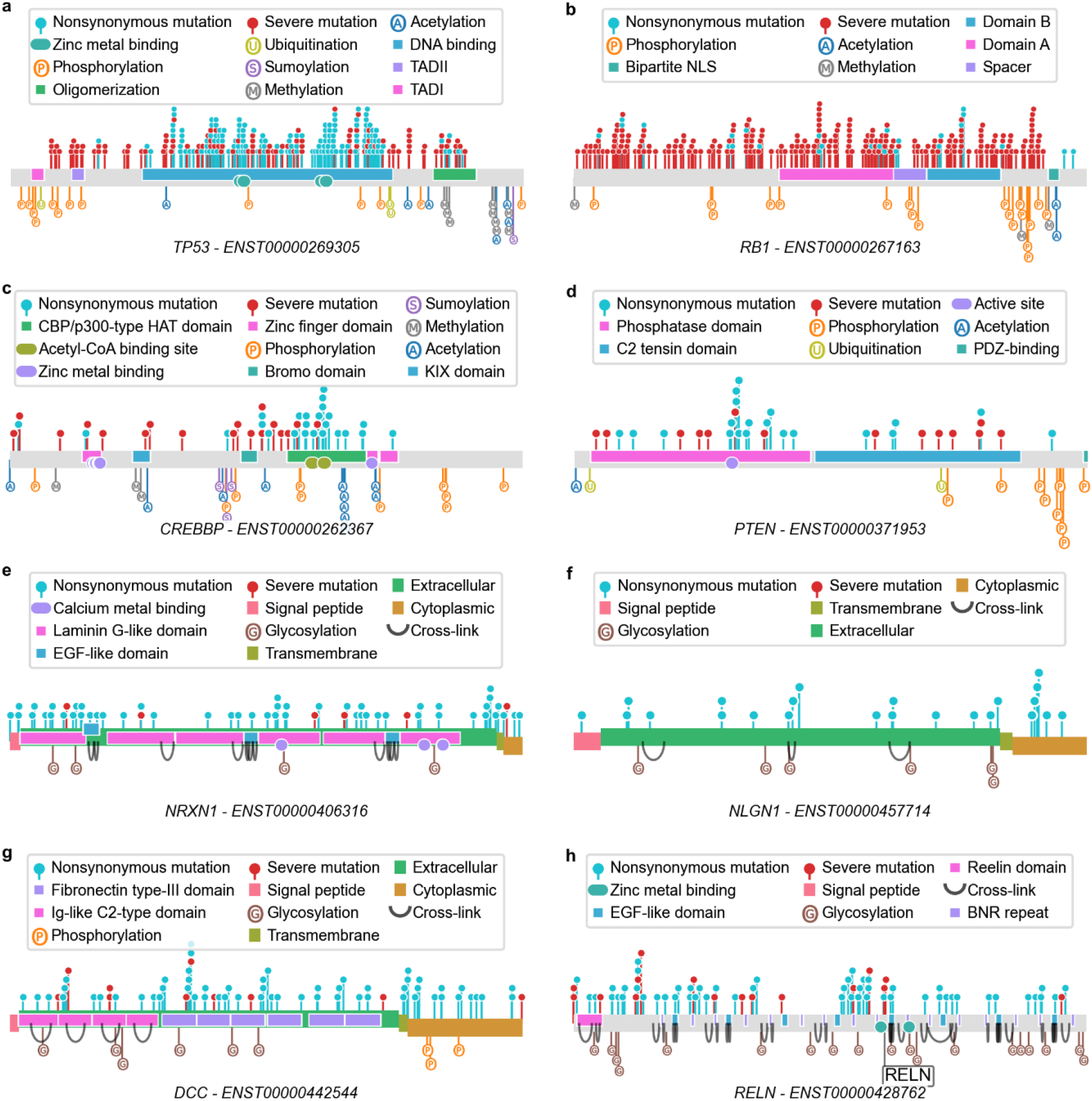
Selected genes mutated in SCLC. Selected genes are shown with the corresponding proteins annotated with UniProt Knowledgebase annotations. Mutations identified in SCLC samples are shown as a lollipop chart above the protein. Severe mutations (stop, frameshift, start-loss, and canonical splice-site mutations) are shown in red. Nonsynonymous mutations (amino-acid substitutions, non-frameshift indels) are shown in light blue. **a)** Mutations in *TP53* are either severe or clustered in the DNA-binding domain. **b)** Mutations in *RB1* are almost exclusively severe. **c)** Mutations in *CREBBP* are severe or clustered in the HAT domain **d)** Mutations in *PTEN* are severe or clustered on the active site. **e)** Mutations in *NRXN1* are mainly nonsynonymous. **f)** Mutations in *NLGN1* are exclusively nonsynonymous **g)** Mutations in *DCC* are mainly nonsynonymous **h)** Mutations in *RELN* are nonsynonymous or severe.

**Extended Data Figure 5:**
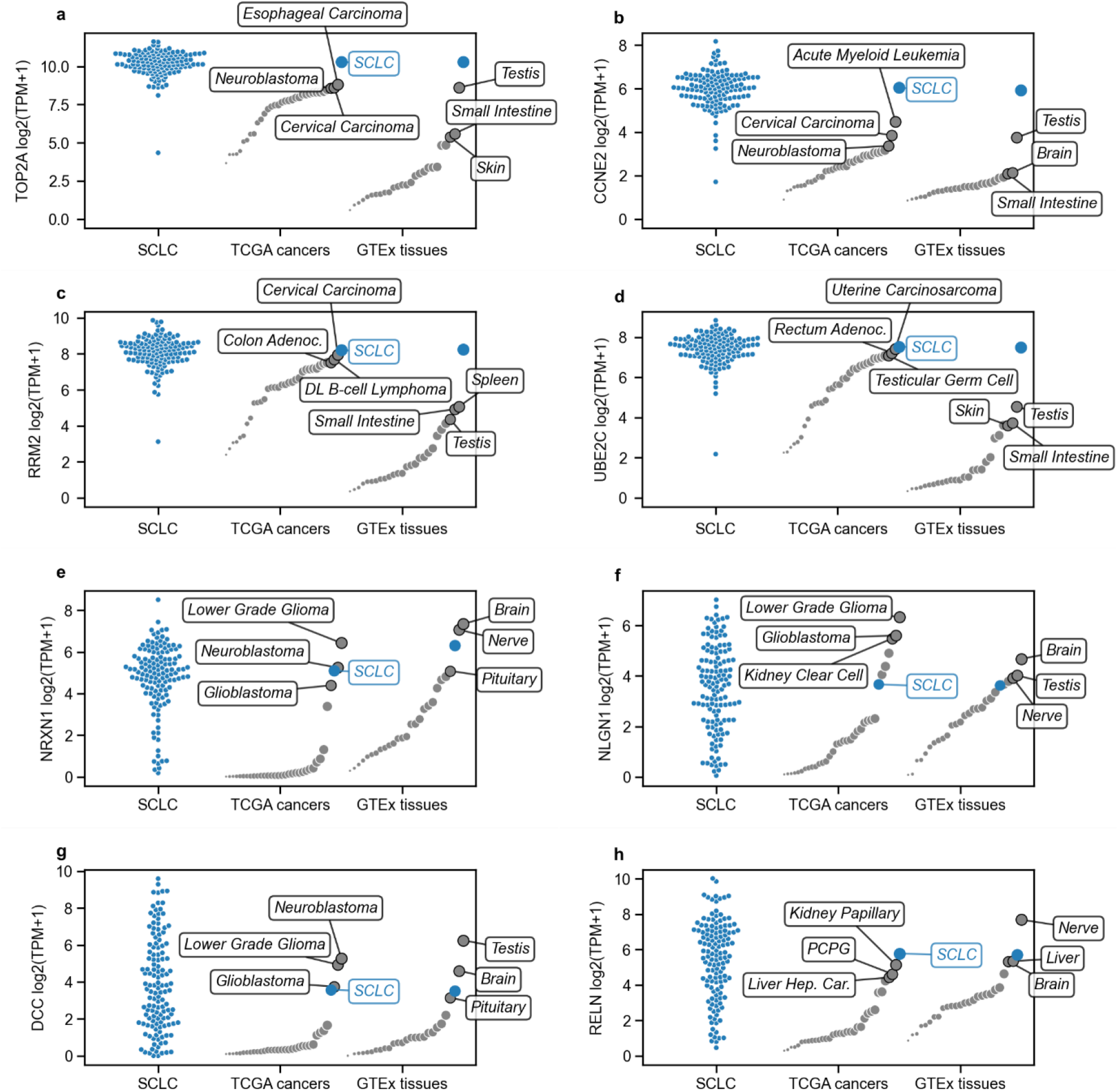
Selected genes specifically expressed in SCLC. The expression levels of individual SCLC samples are shown on the left of each panel. The median expression levels in cancer types included in TCGA and Neuroblastoma as positive control are depicted in the middle. The median expression levels of healthy tissues are on the right. **a, b, c, d)** The expression levels of *TOP2A*, *CCNE2*, *RRM2* and *UBE2C*, exemplary of genes involved in cell-proliferations, are higher in SCLC than in any other cancer or healthy tissue. **e, f, g, h)** The expression levels of *NRXN1, NLGN1, DCC and RELN*, exemplary for synaptic and neuronal genes, are higher in SCLC than in most other cancers and tissues.

**Extended Data Figure 6:**
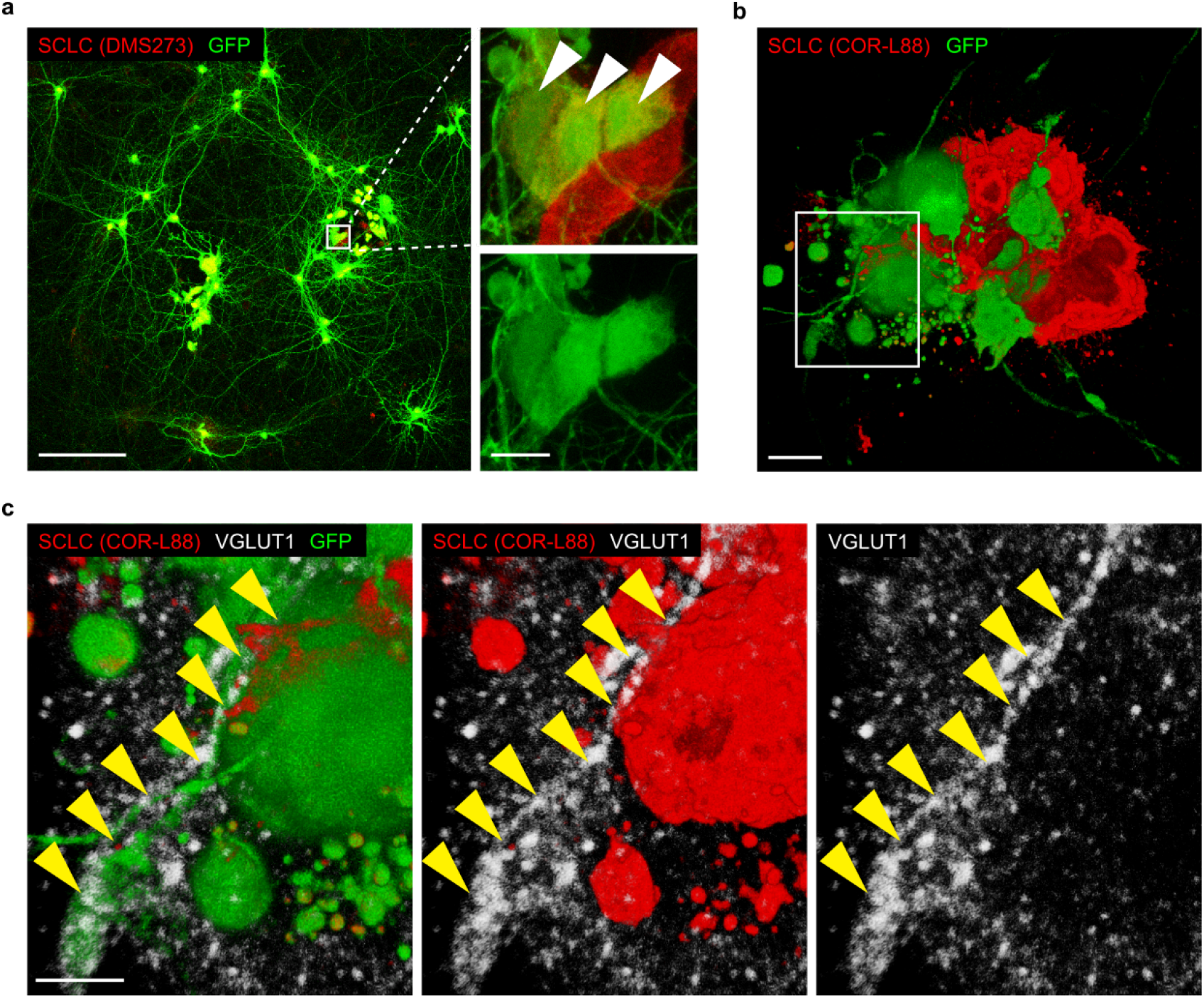
Transmission of rabies virus between SCLC cells and neurons. **a)** RABV-GFP-based tracing of neurons monosynaptically connected to DMS273 SCLC cells expressing DsRed. Right panels show enlarged views of the boxed area containing double-positive starter SCLC cells. Bars, 100 and 10 μm. **b)**3D reconstruction of double-positive starter cells in a cluster of DsRed-expressing SCLC cells (COR-L88) following RABV-GFP-based tracing. Bar, 7 μm. **c)** Magnification of the panel boxed in **b**, showing the profuse expression of VGLUT1-positive punctae in GFP-positive neuronal fibers (yellow arrowheads) contacting starter SCLC cells. Bar, 4 μm.

**Extended Data Figure 7:**
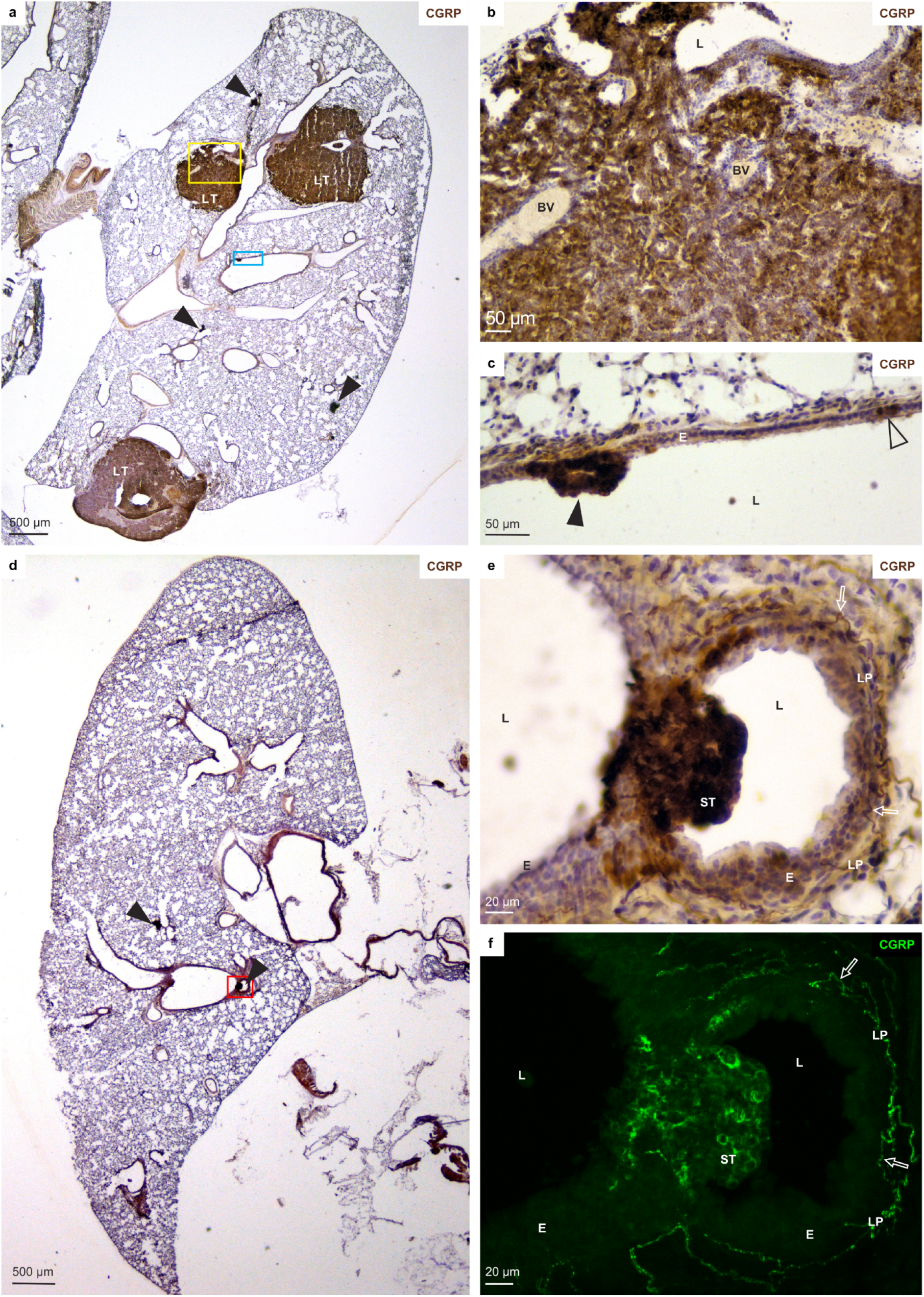
SCLC tumors of different sizes. CGRP immunostaining (brown DAB label) is used to identify the neuroendocrine cells in lung lobes of a *Rb1^fl/fl^;Trp53^fl/fl^* mice. **a)** Tumors were induced eight months before sacrificing the animal. Three large tumors (LT) and several small tumors (arrowheads) are visible. **b)** Magnification of the yellow inset in **a**, showing a gland-like appearance of the tumor. BV: blood vessels. **c)** Magnification of the blue inset in **a**, showing a small proliferation of CGRP+ neuroendocrine cells (arrowhead) and a normal pulmonary NEB (open arrowhead). **d)** Four months after tumor induction. Several small proliferations of neuroendocrine cells (arrowheads) are visible. **e)** Higher magnification of the red inset in **d** showing that the PNECs in a small tumor (ST) bulge into the airway. **f)** Confocal maximum intensity projection of the same section and area as imaged in **e**. FITC-fluorescence reveals the individual CGRP+ PNECs that build up the small tumor and CGRP+ nerve fibers (open arrows) in the lamina propria (LP). L: lumen of airways; E: airway epithelium.

**Extended Data Figure 8:**
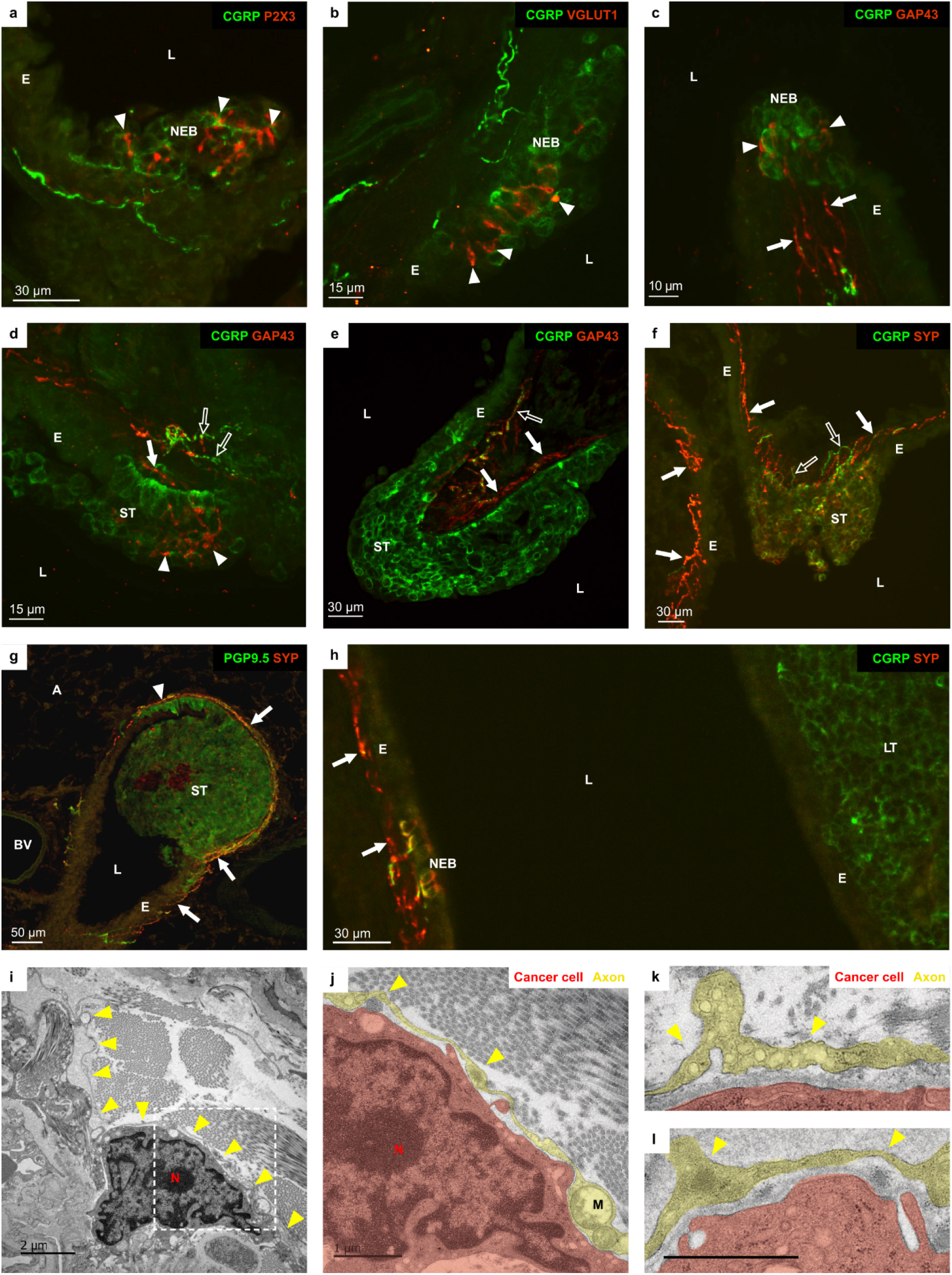
Nerve fibers near murine neuroepithelial bodies and tumors. **a-h)** Lung cryostat sections of *Rb1^fl/fl^;Trp53^fl/fl^* mice. L: lumen of the airway; E: airway epithelium **a)** Intraepithelial P2X3+ (red Cy3 fluorescence) vagal sensory nerve terminals (arrowheads) protruding between the CGRP+ (green FITC fluorescence) neuroendocrine cells of a NEB with normal architecture. **b)** Normal NEB. Intraepithelial VGLUT1+ (red) nerve terminals (arrowheads), representing the glutamatergic vagal sensory subpopulation of NEB innervation, branch between the CGRP+ (green) PNECs. **c)** Branching point of an intrapulmonary airway showing a normal pulmonary NEB. GAP43+ (red) nerve fibers (arrows), as a marker for newly formed and/or remodeling nerve fibers, branch and protrude intraepithelially (arrowheads) between the CGRP+ (green) PNECs.**d)** Small tumor (ST) in an intrapulmonary airway. GAP43+ (red) nerve fibers (arrow) approach the epithelium and branch (arrowheads) between the CGRP+ (green) SCLC cells. CGRP+ nerve fibers (open arrows) are also seen close to the base of the tumor. **e)** CGRP+ (green) small tumor (ST) protruding in an airway. GAP43+ (red) nerve fibers (arrows), a subset of which is CGRP+ (open arrow), are present in the lamina propria but do not protrude between the SCLC cells. **f)** A CGRP+ (green) and SYP+ (red) small tumor (ST) protrudes in an intrapulmonary airway. Abundant SYP+ nerve fibers (arrows) are present in all subepithelial areas. Just beneath the tumor, both SYP+ (arrows) and CGRP+ (open arrows) nerve fibers can be seen. **g)** Intrapulmonary airway and adjacent alveolar (A) region showing a small tumor (ST), double stained for PGP9.5 (green) and SYP (red). The PNECs with a more physiological morphology (arrowhead), likely part of the original NEB from which the SCLC tumor developed, are more intensely stained for PGP9.5. Nerve fibers in the lamina propria (arrows) co-express SYP and PGP9.5. BV: blood vessel. **h)** Intrapulmonary airway with a small part of a large tumor (LT). The SCLC cells show a clear CGRP immunofluorescence (green), while SYP expression (red) appears weak. The epithelium of the same airway at the left side of the image shows a normal pulmonary NEB with CGRP+/SYP+ PNECs and subepithelial SYP+ and CGRP+ nerve terminals (arrows). Remarkable is that the subepithelial area adjacent to the large tumor is devoid of nerve fibers. **i)** Electron micrographs showing a cancer cell surrounded by long axon-like fibers near the periphery of a tumor in the lung of a *Rb1^fl/fl^;Trp53^fl/fl^* mouse. **j,k,l)** Magnifications showing the presence of enlarged structures along identified fibers being compatible with putative boutons containing multiple vesicles and mitochondria (M).

**Extended Data Figure 9:**
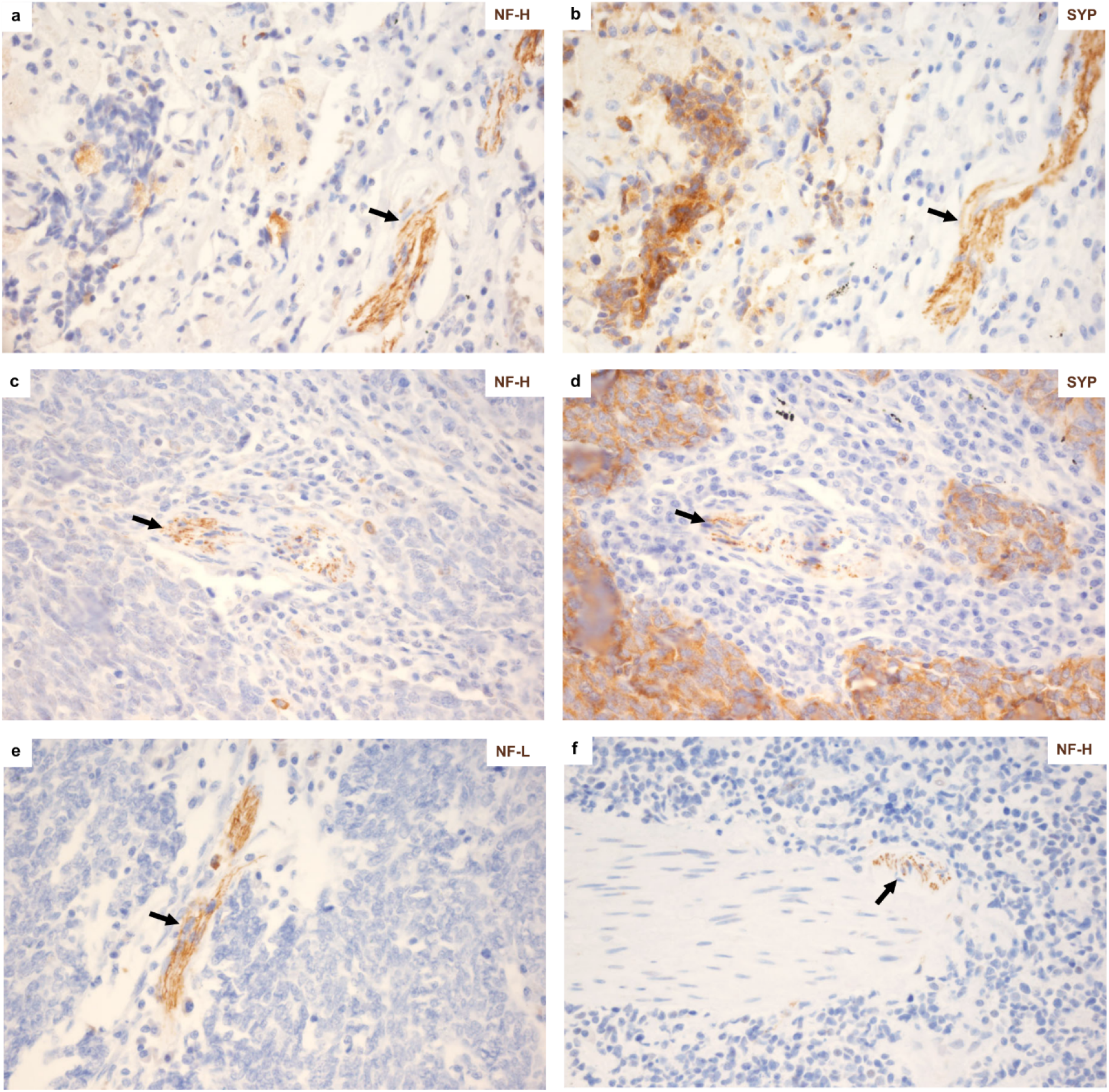
Nerve fibers near SCLC tumors in biopsies from human patients. **a, b)** Neurofilament-heavy polypeptide (NF-H) and synaptophysin (SYP) immunostainings (brown DAB label) show NF-H-positive nerve fibers at the borders of a SYP-positive tumor in a biopsy from a SCLC patient. **c, d)** NF-H and SYP immunostainings (brown DAB label) show NF-H-positive nerve fibers at the borders of a SYP-positive tumor in a biopsy from a second patient. **e)** Neurofilament-light polypeptide (NF-L) immunostaining (brown DAB label) shows a nerve fiber entrapped within the tumor from the second patient **f)** NF-H immunostaining (brown DAB label) show NF-H-positive nerve fibers near an intratumoral vessel in the biopsy from a third patient. All sections are counterstained with hemalum. Original microscopic magnifications were 400x.

**Extended Data Figure 10:**
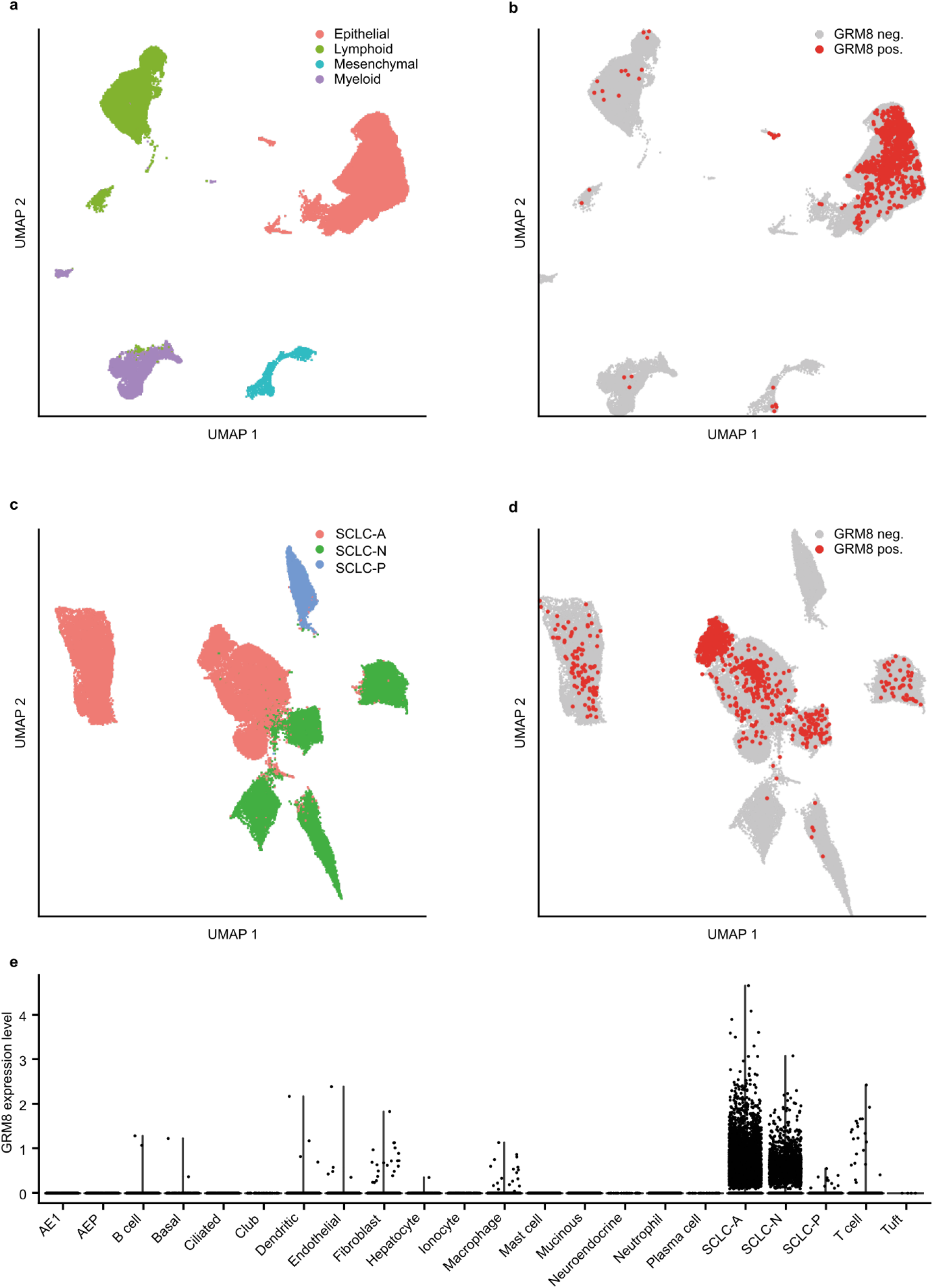
*GRM8* is specifically expressed in cancer cells. **a)** Uniform Manifold Approximation and Projection (UMAP) of scRNA-seq derived from 21 SCLC and 4 normal human lung samples (n = 89238 cells) from Chan *et al*. ^44^. Colors indicate cell lineage annotation. **b)** Same projection as in **(a)** highlighting cells with detected *GRM8* expression (red) compared to *GRM8* negative cells (grey). **c)** UMAP projection of only SCLC cells (n=54313). Colors indicate SCLC subtype as *ASCL1* positive (SCLC-A, orange), *NEUROD1* positive (SCLC-N, green) or *POU2F3* positive (SCLC-P, blue). **d)** Same projection as in **(c)** highlighting cells with detected *GRM8* expression (red) compared to *GRM8* negative cells (grey). **e)** Detailed analysis of *GRM8* expression level across fine-grained cell type classification in cells of SCLC and normal samples displayed in **(a)**.

## Materials and Methods

### Mice

This study was performed in accordance with FELASA recommendations, with European Union guidelines and German guidelines. The experiments were approved by the local Ethics Committee of Animal experiments (Landesamt für Natur, Umwelt und Verbraucherschutz Nordrhein-Westfalen). The mice were housed in groups of up to five animals per cage supplied with standard pellet food and water *ad libitum* with a 12 h light/dark cycle, while temperature was controlled to 21-22°C.

### SCLC tumor induction

In order to induce lung tumor formation and, when present, activation of the *piggyBac* transposition system, eight-to twelve-week-old mice were anesthetized with Ketavet (100 mg/kg) and Rompun (20 mg/kg) by intraperitoneal injection followed by intratracheal instillation of replication-deficient adenovirus expressing Cre-recombinase (Adeno-Cre, 2.5 x 10^7^ PFU). Viral vectors were provided by the University of Iowa Viral Vector Core (http://www.medicine.uiowa.edu/vectorcore).

### MR imaging

Mice were anesthetized with 2.5% isoflurane. An Achieva 3.0 T clinical MR imaging system (Philips Healthcare, Best, the Netherlands) in combination with a dedicated mouse solenoid coil (Philips Healthcare, Hamburg, Germany) were used for imaging. Animals were anaesthetized using isoflurane (2.5%) and T2-weighted MR images were acquired in the axial plane using a turbo-spin echo (TSE) sequence (repetition time [TR] = 3819 ms, echo time [TE] = 60 ms, field of view [FOV] = 40 × 40 × 20 mm3, reconstructed voxel size = 0.13 × 0.13 × 1.0 mm3, number of average =1). MR images (DICOM files) were analyzed by determining and calculating region of interests (ROIs) using the Horos software.

### *PiggyBac* transposition system in SCLC

For the activation of transposition in a SCLC mouse model, we used the following alleles, as detailed in **Fig. S1**: *Rosa26^LSL-PB^, ATP1-S2, ATP1-H39, Rb1^flox^* and *Trp53^flox^* ^17,18^. The mice were kept on a mixed *C57Bl6/Sv129* background. Genotyping primers are reported in the **Extended Data Table 9**. The ATP1 alleles were genotyped using ATP-F and ATP-R primers. The *Rosa26^LSL-PB^* knock-in allele was genotyped using BpA5F and Rosa3R primers. The wild-type *Rosa26* allele was detected with the Rosa5F and Rosa3R primers. To study SCLC formation, all four mouse lines were aged following adenoviral instillation, as described above. After reaching the termination criteria, mice were sacrificed and single tumor nodules were isolated and used for DNA extraction. Analysis of transposon mobilization at the donor locus and splinkerette-PCR amplification of transposon insertion sites were performed as previously described ^18,51^.

### Treatment of *piggyBac* mice

Starting five months after tumor induction, tumor growth was monitored by bi-weekly MR imaging as described above until termination criteria were reached. Upon tumor detection (minimal tumor size 3mm^3^), RPLS and RPLH mice were treated with either a combination of cisplatin and etoposide or anti-PD1 antibody RPM1-14. Compound solutions were prepared and injected as follows: Etoposide (Hexal) was administered on days 1,2 and 3 of a 14-day cycle, i.p., at a concentration of 10 mg/kg. Cisplatin (Accord) was administered on day 1 of 14-days cycle at a concentration of 5 mg/kg, i.p. The anti–PD-1 antibody RMP1-14 (BioXCell) was administered i.p. 2 days per week (250 μg/administration).

### Reference genomes and gene definitions

The reference genome used for all human analyses was the TCGA GRCh38.d1.vd1, with the exception of the comparison of human RNAseq data to the GTEx data, which was performed using the GTEx v8 reference (Homo_sapiens_assembly38_noALT_noHLA_noDecoy_ERCC.fasta). The reference genome used for all mouse analyses was the Ensembl version GRCm38 (Mus_musculus.GRCm38.dna.fa). The gene annotation for analyses of human genetic data was the gencode annotation v22, while the gene annotation for analyses of mouse data was the gencode annotation vM23 ^52^. Both gencode annotations were filtered first to only include transcripts marked as “protein coding” and subsequently to only include the 17153 genes for which a one-to-one ortholog could be identified between mouse and human using the HCOP fifteen-column orthology table (downloaded on the 06.01.2020 from the HGNC database ^53^). The gene annotation for the analysis of TCGA expression data was the gencode annotation v22 and the gene annotation for the analysis of GTEx expression data was the gencode annotation v26.

### Analysis of *piggyBac* insertions

Sequencing reads that contained internal transposon sequences were excluded and the remaining reads were aligned against the GRCm38 reference. The aligned reads that did not align to the consensus TTAA target sequence were excluded. At each TTAA locus in each sample, reads derived from the same fragment, identified by the identical position of the read ends, were collapsed. TTAA loci were kept if five or more different fragments were identified. Germline insertions were identified by the presence of 10 or more different fragments at a TTAA locus in the tail or ear samples. These TTAA loci were excluded from analysis in the whole cohort and 1 Megabase upstream and downstream were masked from analysis of tumors from the affected mice. 10 Megabases around the donor loci were also masked from analysis (chromosome 5:50000000-70000000 for the RPLH line and chromosome 10:0-10000000 for the RPLS line). Insertions detected in more than one tumor were assigned to the tumor with the highest number of fragments. For each of the 17153 protein coding genes present in both the human and mouse genomes we defined the included genomic range as the union of all the transcripts of the gene from the transcription start site to the stop codon. The statistical analysis included two steps. First, at the sample level, the Poisson distribution was used to calculate the probability of seeing at least as many transposon fragments as actually present. The rate used for the Poisson distribution was based on the total insertion rate within genes of each chromosome of each sample, on the total number of TTAA sites within genes of the chromosome and on the number of TTAA sites within each gene. We then calculated FDR-corrected q values for each sample and each gene. We obtain a total of 11208 genes (an average of 37 genes per sample) which are significant at a cutoff of q < 0.05 at the sample level. To calculate the statistical significance of the genes at the cohort level, we again used the Poisson distribution with a rate derived from distributing the 11208 hits evenly across all non-masked genes of all samples. We then calculate the FDR-corrected q values at the cohort level for each gene.

### Analysis of *PiggyBac* sub-cohorts

We used a permutation test to compare the distribution of the transposon insertions in different sub-cohorts. For each comparison, the union of samples included in the comparison was shuffled 100000 times, while maintaining the same number of samples from each mouse line in each sub-cohort (RPLH and RPLS). For each gene, we then counted the number of iterations in which the absolute difference in the fraction of samples carrying an insertion is greater than in the real configuration. We then calculated the FDR-corrected q values for each gene.

### Simulation and annotation of possible human mutations

For each gene included in the filtered gencode annotation v22, all possible single-nucleotide substitutions were simulated, annotated using Annovar v2018Apr16 ^54^ with the filtered gencode annotation v22, and divided into three categories: synonymous (no predicted change in the protein sequence), severe (causing a premature stop, the loss of the starting ATG site, a frameshift or a nucleotide change in one of the two intronic bases flanking each side of an exon) and nonsynonymous (any other predicted change in the protein sequence). For each simulated variant on each gene, only the most severe consequence among all the transcripts associated with the gene was kept. All simulated variants were also annotated using the total population frequency in non-cancer samples from the gnomAD v2.1.1 GRCh38 liftover exome and the gnomAD v3 genomes and excluded if they are found in more than 1 in 10000 samples. Based on this simulation, the number of possible nonsynonymous or severe variants for each gene was used as the basis for the calculation of the expected number of mutations in each gene.

### Data collection of human somatic mutations

Sample information and mutations were downloaded from the supplementary tables of the respective papers or from the CCLE website (Cell_lines_annotations_20181226.txt and CCLE_DepMap_18q3_maf_20180718.txt, https://portals.broadinstitute.org/ccle/). Where needed, the mutations were mapped to the TCGA GRCh38 reference (GRCh38.d1.vd1.fa) using the liftOver tool from the USCS database (http://genome.ucsc.edu, ^55^) The resulting 177983 mutations were annotated as described above for the simulated variants. 613 mutations were excluded from analysis (517 mapped to mitochondrial genes and 96 could not be mapped to primary chromosomes in h38). The remaining 177370 variants were left-aligned using GATK LeftAlignAndTrimVariants v4.1.3.0 ^56^. A total of 28 samples were excluded from analysis because they shared 5 or more mutations with a sample from a more recent study, leaving 456 samples.

### Analysis of human sub-cohorts

We used Fisher’s exact test to compare the distribution of the mutations within different sub-cohorts: primary versus metastatic samples, treated versus untreated samples and cell lines versus tumor samples. For each gene, the contingency table was composed of the number of mutations without and within the gene for each subcohort. The resulting p values were corrected for multiple testing with the false discovery rate.

### Statistical analysis of the human cohort

Samples sharing five or more mutations were merged (eg, samples sequenced both before and after treatment). In total, 439 samples and 117353 nonsynonymous mutations were used for analysis. We used the Poisson distribution to estimate the chances of observing at least as many mutations by chance in each gene. To obtain the rate for the Poisson distribution for each gene, we divided the total number of nonsynonymous mutations, counting each sample at most twice per gene, by the total number of possible nonsynonymous mutations within the 17153 protein coding genes present in both the human and mouse genomes. For each gene, we then multiplied this value by the number nonsynonymous mutations that are theoretically possible in the gene (see simulation above). The rate therefore represented the expected number of nonsynonymous mutations under a uniform distribution model. For each gene we then calculated the probability of observing at least as many mutations as actually present. We corrected the resulting p values for multiple testing with the false discovery rate to derive the q-value for each gene. Finally, we repeated this analysis but only included severe mutations (stop-gain, start-loss, frameshift and canonical splicing) to derive the probability of observing at least as many severe mutations as actually present. Mutations in selected genes were plotted on the corresponding proteins with annotations derived from the Uniprot Knowledgebase (https://www.uniprot.org/, accessed on June 14^th^ 2022) ^57^.

### Comparison of expression data to the TCGA database

SCLC RNAseq data from two different studies ^2,35^ and RNAseq data from neuroblastoma samples ^58^ were reanalyzed using the TCGA pipeline. Briefly, STAR version 2.4.2a was used to align the reads to the GRCh38 reference using the gencode annotation version 22. HTSeq version 0.6.1p1 was then used to quantify the expression at the gene level. The raw counts were converted to TPM using the median length of all transcripts for each gene, as reported in the gencode annotation version 22. The TPM +1 values were then logscaled and used for further analysis. Expression data were downloaded from the Genomic Data Commons Data Portal (https://portal.gdc.cancer.gov). The TPM values of SCLC samples were compared to the TPM values of the individual types of tumors within TCGA using a two-sided Mann-Whitney test and the fold change for each gene was calculated as the median of the SCLC log2(TPM+1) values minus the median of the TCGA cohort log2(TPM+1) values.

### Comparison of expression data to the GTEx database

SCLC RNAseq data from two different studies ^2,35^ were reanalyzed using the GTEx pipeline v8. Briefly, STAR version 2.5.3a was used to align the reads to the GRCh38 reference using the gencode annotation version 26. RNA-SeQC version 1.1.9 was then used to quantify the expression at the gene level. The raw counts were converted to TPM using the median length of all transcripts for each gene, as reported in the gencode annotation version 26. The TPM +1 values were then logscaled and used for further analysis. Expression data were downloaded from the GTEx database (https://gtexportal.org). Tissues with less than 30 samples were excluded (Fallopian Tube, Bladder, Cervix Uteri). The TPM values of SCLC samples were compared to the TPM values of the individual tissues within GTEx using a two-sided Mann-Whitney test and the fold change for each gene was calculated as the median of the SCLC log2(TPM+1) values minus the median of the GTEx tissue log2(TPM+1) values.

### Gene set analysis with the Gene Ontology

The Gene Ontology (GO) architecture and annotations were downloaded from the GO website (http://geneontology.org, accessed on September 13^th^ 2020) ^36,37^. For each term annotation of each gene, the gene was additionally annotated with all its parent terms. For each dataset of interest, the identified genes were compared to all GO terms that included at least 10 and at most 1000 genes with Fisher’s exact test and FDR correction. *piggyBac* hits (n=504) and human mutation hits (n=991) were included if they had a q value < 0.1 and at least one GO annotation. Genes highly expressed in SCLC were first filtered to only include genes with at least one GO annotation and with q values less than 0.1 in at least 90% of the comparisons (30/33 tumors or 25/27 healthy tissues). The remaining genes were then ranked by the median log2 fold change across all comparisons and only the top 1000 genes were included. The force-directed graphs were generated with datashader v0.12.1 using the ForceAtlas2 layout. For this analysis, up to 100 GO terms were included as nodes if they were significantly enriched (q < 0.1) for genes in the datasets and if they were not a perfect subset or overset of a GO term with a more significant overlap. An edge was present between two GO terms if the genes included in the terms were significantly overlapping (q<0.1 after Fisher’s exact test and FDR correction). Go terms identified at the expression level both versus cancers and versus healthy tissues were further cross-referenced with Chip-seq data downloaded from the CISTROME database (http://cistrome.org/db, accessed November 27^th^ 2019) ^39^ and with scRNA-seq data from healthy human lung from Travaglini *et al*.^40^. The CHIPseq peaks from experiments using antibodies against RB1, RBL2, E2F1, E2F2, E2F3, E2F4, E2F5 were downloaded from the CISTROME database to derive an RB-E2F score. For each gene, the peaks were merged across samples and replicates and their fold change over the background was added across samples and replicates. Peaks with a total fold change of at least 10 were included in the analysis. The regulatory potential was calculated for all target genes whose transcription start sites (TSS) was within 100 kb of the peak, using the distance between the peak and the TSS, as described by Tang et al. ^59^. The regulatory potential was multiplied by the total fold change and this score was added for all peaks near a TSS. For each target gene, the transcript with the highest score was kept. The scores for each transcription factor were normalized between 0 and 1 and then added together to derive the RB-E2F score for each target gene. The score for each enriched GO term was then calculated as the mean score across genes included in the GO term and in the SCLC dataset. ScRNA-seq data, as well as the corresponding metadata from Travaglini et al. was downloaded from Synapse (Synapse:syn21560406). Cell type annotations were obtained from the metadata. The cells were divided in two groups: cells annotated as Neuroendocrine and all others. UMIs within each group were added and converted to TPM. The TPM+1 values were then log scaled. The log2 fold change between PNECs and other lung resident cells was then calculated as the difference of the two log scaled values for each gene. For each GO term, we then calculate the mean fold change of genes included in the GO term and in the SCLC dataset.

### Neuroendocrine and *Glutamatergic Synapse* expression scores

To calculate the neuroendocrine expression score, the log-scaled expression values of the neuroendocrine markers *SYP, CHGA, ENO2, NCAM1* and *S100B* were first normalized between the median of the tumor type or tissue with the highest expression (normalized to 1) and the median of the tumor type or tissue with the lowest expression (normalized to 0). The score was then calculated as the mean of the normalized expression for the five markers. The same process was used to calculate the expression score for genes within the GO term *Glutamatergic Synapse*.

### Reanalysis of scRNA-seq data from SCLC patients

Published scRNA-seq data ^44^ were obtained from https://cellxgene.cziscience.com/collections containing pre-processed gene expression values, annotations of cell types, SCLC subtype and Uniform Manifold Approximation Projection (UMAP) embeddings. Distribution of cells from 21 SCLC and 4 normal lung samples were visualized using UMAP projections followed by addition of information on cell types or *GRM8* expression based on normalized expression > 1. In addition, *GRM8* expression was quantitatively assessed using violin plots per cell type detected in the 25 samples.

### Viral production

Retroviruses encoding for *DsRedExpress2* and for the RABV glycoprotein (*G*) and the *TVA800* (the GPI anchored form of the TVA receptor), as well as the GFP-encoding EnvA-pseudotyped RABV were described previously ^60^.

### Cell culture

Mouse embryos (E13.5) were isolated following cervical dislocation of the anesthetized pregnant mother as previously described ^61^. Briefly, cortices were dissected in Hank’s buffered salt solution (HBSS, Gibco) supplemented with HEPES (10 mM, Gibco), and dissociated by means of enzymatic digestion for 20 min at 37°C by incubating the tissue in DMEM high Glucose Glutamax (Gibco) containing papain (20 U/ml, Merck) and cystein (1 μg/ml, Merck), followed by mechanical trituration in medium supplemented with 10% fetal bovine serum (Gibco). Cells were plated at a density of 0.4 × 10^5^ cells/cm^2^ on poly-L-lysine (0.1 mg/ml, Merck) coated glass coverslips. After four hours, the medium was replaced with Neurobasal serum-free medium (Gibco) containing B27 supplement (1%, Gibco) and Glutamax (0.5 mM, Merck). Neurons were then maintained at 37°C and 5% CO2 throughout the experiment and semi-feeding was performed once per week. COR-L88, DMS273, H146, H69 and H524 human SCLC cell lines were maintained in culture in RPMI 1640 (Life Technologies) supplemented in 10% fetal bovine serum (Gibco) and 1% Pen/Strep (Gibco). For experiments requiring co-cultures of neurons and SCLC cells, neurons grown for at least seven days in vitro (DIV7) were used. SCLC cells preconditioned in neurobasal medium for 48h were plated at a density of 1000 cells onto the neuronal layer. As a control, monocultures of SCLC cells maintained in neurobasal medium were used.

### RABV tracing

Neuronal cultures added with SCLC cells which had been previously transduced with a G-TVA-DsRed retrovirus were used. At neuronal DIV10-12, the EnvA-pseudotyped RABV encoding for GFP was added to the medium. Analysis was then performed after additional 2-4 days in culture, when samples were fixed in PFA (4% in PBS; Sigma) and the number of GFP-positive only presynaptic neurons quantified and normalized to the number of double-positive starter SCLC cells. The p values were calculated using a one-sided Mann-Whitney test.

### Proliferation assays

To compare proliferation rates of mono-cultures and co-cultures with immunofluorescence stainings, cultures of COR-L88 cells were treated with 20μM EdU and incubated for 2h prior to fixation with 4% PFA pre-warmed at room temperature for 5 min. Cells were washed three times with 1X PBS followed by EdU staining using the Click-iT EdU Imaging Kit (Invitrogen). The p values were calculated with a one-sided Mann-Whitney test. For the calculation of the population doublings with and without neurons, SCLC cell lines (COR-L88, DMS273, H146, H69, H524) were pre-stained with the live cell labelling kit Cytopainter (ab187964) according to the manufacturer instruction, and seeded either alone or with neurons at day DIV9. After three and six days, cells were collected, washed with FACS buffer (PBS, 3% FBS) and stained DAPI prior to counting with a MACSQuant analyser-16 flow cytometer and analysis with Floreada.io legacy version. The p value was calculated with a two-sided Wilcoxon rank sum test and adjusted for false discovery rate.

### Transplantation experiments

Thy1-GFP-M mice ^62^ were anesthetized by intraperitoneal injection of a ketamine/xylazine mixture (100 mg/kg body weight ketamine, 10 mg/kg body weight xylazine), treated subcutaneously with Carprofen (5 mg/kg) and fixed in a stereotactic frame provided with a heating pad. A portion of the skull covering the somatosensory cortex (from Bregma: caudal: −2.0; lateral: 1.5) was thinned with a dental drill avoiding to disturb the underlying vasculature and small craniotomy sufficient to allow penetration of a glass capillary was performed. A finely pulled glass capillary containing the suspension of murine SCLC cells derived from the *Rb1^fl/fl^;Trp53^fl/fl^* model in sterile PBS was then inserted through the dura to reach the hippocampus (−1.9 to −1.3 from Bregma) and an estimated total of about 30-50.000 cells (corresponding to a total injected volume of 0.8-1.0 μl) were slowly infused via a manual syringe (Narishige) in multiple vertical steps spaced by 50 μm each during 10-20 minutes. After capillary removal, the scalp was sutured and mice were placed on a warm heating pad until full recovery. Physical conditions of the animals were monitored daily to improve their welfare before euthanize them.

### Fluorescence immunostaining of brain slices and co-cultures

Immunostaining of fixed brain slices and cultures was performed using conventional procedures described previously ^61^. The following primary antibodies were used: chicken anti-GFP (1:500, Aves Labs, GFP-1020), rabbit anti-RFP (1:500, Rockland, #600401379), chicken anti-MAP2 (1:500, Abcam, #ab5392), mouse anti-VGlut1 (1:500, Synaptic Systems, #135311). The following secondary antibodies were used (raised in donkey): Alexa Fluor 488-, Alexa Fluor 546-, Alexa Fluor 647-conjugated secondary antibodies to rabbit, mouse, chicken (1:1000, Jackson ImmunoResearch). Images were acquired utilizing an SP8 confocal microscope (Leica) equipped with a 20x (NA 0.75), 40x (NA 1.3), 63x (NA 1.4) or 100x (NA 1.3) oil immersion objective and further processed with Fiji.

### Fluorescence immunostaining of lung cryostat sections

*Rb1^fl/fl^;Trp53^fl/fl^* mice were sacrificed four and eight months after tumor induction. Lungs were fixed by intratracheal instillation with 4% paraformaldehyde in 0,1M phosphate buffer and processed to collect cryostat sections ^16^. Immunostaining was performed as previously described for mouse lungs ^16^. The following primary antibodies were used: goat anti-CGRP (1:1000, Abcam, #ab36001), rabbit anti-GAP43 (1:2000, Novus Biologicals, #NB300-143), rabbit anti-PGP9.5 (1:2000, Abcam, #ab108986), rabbit anti-P2X3 (1:1000, Chemicon, #AB5895), guinea-pig anti-SYP (1:4000; Synaptic Systems, #101002), rabbit anti-VGLUT1 (1:250, Synaptic Systems, #135303). The following secondary antibodies were used (raised in donkey): FITC- or Cy3-conjugated, secondary antibodies to rabbit, goat and guineapig (1/200-1/2000; Jackson ImmunoResearch). To enhance staining intensity or visualize sections in a conventional light microscope with DAB, biotinylated secondary antibodies (1:500, Jackson ImmunoResearch) were combined with Cy3-conjugated streptavidin (1:6000; Jackson ImmunoResearch) or ExtrAvidin-HRP (1:1000, Sigma,#E2886), respectively. Confocal images were acquired using a microlens-enhanced dual spinning disk confocal microscope (UltraVIEW VOX, PerkinElmer) equipped with 488 nm and 561 nm diode lasers for excitation of respectively FITC and Cy3.

### Immunohistochemistry

Patients consented to the use of their tissue specimens and approval was obtained by the Ethics Committee of the University of Cologne (Biomasota # 13-091, 2016). Formalin-fixed and paraffin-embedded tissue sections of human SCLC tumors were deparaffinized and immunohistochemically stained according to standard protocols using an automated immunostainer and a horseradish peroxidase-based detection system with diaminobenzidine as chromogen. Primary mouse monoclonal antibodies were directed against synaptophysin (1:100, Leica Biosystems, Wetzlar, Germany, #PA0299), neurofilament, 200 kDa subunit (NF-H) (1:500, Sigma, Saint Louis, MO, #N0142), and neurofilament, 70 kDa subunit (NF-L, 1:500, Agilent, Santa Clara, CA, #M0762). Immunostained sections were counterstained with hemalum.

### Transmission Electron Microscopy

Anesthetized mice were transcardially perfused with a fixative solution containing 4% formaldehyde and 2.5% glutaraldehyde in 0.1 M cacodylate buffer. The lungs were isolated, cut in 1 mm thick sagittal sections and the examined area was dissected according to the location of the tumor mass. EPON embedding and ultrathin sections were prepared using standard procedures ^63^. Electron micrographs were taken with a JEM-2100 Plus Transmission Electron Microscope (JEOL), equipped with Camera OneView 4 K 16 bit (Gatan) and software DigitalMicrograph (Gatan). For analysis, electron micrographs were acquired with a digital zoom of 5000x or 6000x.

### Electrophysiology

Acutely isolated brains were sectioned into coronal slices of 300 μm by using a vibrating microtome (HM-650 V, Thermo Fisher Scientific, Walldorf, Germany) filled with ice-cold, carbogenated (95% O2 and 5% CO2) aCSF cutting solution (in mM: 125.0 NaCl, 2.5 KCl, 1.25 sodium phosphate buffer, 25.0 NaHCO3, 25.0 D-glucose, 1.0 CaCl2, and 6.0 MgCl2, adjusted to pH 7.4 and 310 to 330 mOsm). The obtained brain slices were transferred into a chamber containing aCSF recording solution (in mM: 125.0 NaCl, 2.5 KCl, 1.25 sodium phosphate buffer, 25.0 NaHCO3, 25.0 D-glucose, 4.0 CaCl2, and 3.5 MgCl2, adjusted to pH 7.4 and 310 to 320 mOsm). Slices were stored for at least 30 min to allow recovery before performing recordings.

All recordings were performed using a microscope stage equipped with a fixed recording chamber and a 20× water-immersion objective (Scientifica). For *ex vivo* experiments recordings were performed in aCSF recording solution. For *in vitro* experiments SCLC cells from day 3 - 6 in co-cultures were used, and recordings were performed in extracellular solution (in mM: 124 NaCl, 10 D-glucose, 10 HEPES-KOH (pH 7.3), 3 KCl, 2 CaCl2, 1 MgCl2, adjusted to pH 7.4). Patch pipettes with a tip resistance of 5 to 10 megohms were made from borosilicate glass capillary tubing (GB150-10, 0.86 mm × 1.5 mm × 100 mm, Science Products, Hofheim, Germany) with a horizontal pipette puller (P-1000, Sutter Instruments, Novato, CA). The patch pipette was filled with internal solution (in mM: 4.0 KCl, 2.0 NaCl, 0.2 EGTA, 135.0 K-gluconate, 10.0 Hepes, 4.0 ATP(Mg), 0.5 guanosine triphosphate (GTP) (Na), and 10.0 phosphocreatin, adjusted to pH 7.25 and osmolarity 290 mOsm (sucrose). Recordings were performed with an ELC-03XS npi patch-clamp amplifier (npi electronic GmbH, Tamm, Germany) that was controlled by the software Signal (version 6.0, Cambridge Electronic, Cambridge, UK). The experiments were recorded with a sampling rate of 12.5 kHz. The signal was filtered with two shortpass Bessel filters that had cutoff frequencies of 1.3 and 10 kHz. Capacitance of the membrane and pipette was compensated by using the compensation circuit of the amplifier. All experiments were performed under visual control using an Orca-Flash 4.0 camera (Hamamatsu, Geldern, Germany) controlled by the software Hokawo (version 2.8, Hamamatsu, Geldern, Germany).

SCLC cells were identified by the expression of the cytosolic fluorescent protein DsRed. Cells were clamped at a holding potential of −30 mV after rupturing the membrane and spontaneous activity was recorded for 5 min in whole-cell voltage clamp mode. For *ex vivo* experiments, synaptic inputs were isolated by adding the following synaptic blockers to the recording aCSF solution: cyanquixaline (CNQX, sigma C127, 10mM), 2-Amino-5-phosphonopentanoic acid (AP5, sigma A8054, 20μM) and Bicuculline (Bicuculline-Methiodid, sigma 14343, 100μM). For *in vitro* experiments, synaptic inputs were isolated by adding the following blockers to the extracellular solution: riluzole (riluzole, Sigma, 10μM), cyanquixaline (CNQX, sigma, 10mM) and 2-Amino-5-phosphonopentanoic acid (AP5, sigma, 25μM). Spontaneous post-synaptic currents were identified and measured with Igor Pro (version 32 7.01, WaveMetrics, Lake Oswego, OR, USA) using a semiautomatic identification script. The p values were calculated with a one-sided Mann-Whitney test.

### Preclinical SCLC mouse model

For preclinical experiments, we used the *Rb1^fl/fl^;Trp53^fl/fl^* genetically engineered mouse model for SCLC, which is driven by Cre-inducible conditional *Rb1* and *Trp53* knockout, as previously described ^17^. Tumors were induced and monitored via MRI as described above. Upon tumor detection (minimal tumor size 5mm^3^), the mice were randomly distributed into the treatment cohorts. Compound solutions were prepared and injected as follows: Etoposide (Hexal) was administered on days 1,2 and 3 of a 14-day cycle, i.p., at a concentration of 8 mg/kg. Cisplatin (Accord) was administered on day 1 of 14-days cycle at a concentration of 4 mg/kg, i.p. Riluzole (15 mg/kg) was dissolved in 10% DMSO, 40% PEG300, 5% Tween-80, 45% PBS and administered 5 days per week. DCPG (60 mg/kg) was dissolved in PBS and administered i.p. for 5 days per week. Best response was calculated as the lowest percent change measured from the last MRI scan before treatment. The q values for the best response were calculated with Mann-Whitney test followed by FDR correction. The q values for the survival were calculated with the log-rank test followed by FDR correction.

## Data availability

External data are available from the original sources, as described in the methods. The remaining data that support the findings of this study, including source data for the figures, are available from the corresponding authors upon reasonable request.

## Code availability

All code generated during this study is available from the corresponding authors upon reasonable request.

## Materials & Correspondence

Correspondence and requests for materials should be addressed to Christian Reinhardt or Filippo Beleggia

## Author contributions

M.B., E.M., H.C.R., F.B. designed the study. K.C., R.D.J., J.G, S.v.K., M.P., T.P., H.G., M.L.S., J.B, G.R., M.F., D.A., R.B., I.B., R.R., R.K.T, M.B, E.M., H.C.R., F.B. designed the experiments. A.S., V.S., K.N., G.A.W., M.T., I.P., I.K., O.I., J.W., R.M., C.M.B., J.G, M.J., F.O., A.P., A.A.H., M.B., A.H., I.B., M.B., E.M., F.B. performed the experiments. K.N., G.S.G., J.B., G.R., D.A., I.B., M.B., E.M., F.B. performed data analysis. A.S., K.N., I.P., J.B., G.R., I.B., M.B., E.M., F.B. generated the figures. M.B., E.M., H.C.R., F.B. wrote the manuscript.

## Acknowledgements

We are indebted to our patients, who provided primary material. We thank Bianca Göbel, Jacqueline Knufer, Alexandra Florin, Marion Müller, Ursula Rommerscheidt-Fuß from the University Hospital Cologne, for their outstanding technical support and the CECAD Imaging Facility in Cologne for the use of the electron and light microscopes. This work was funded through the German-Israeli Foundation for Research and Development (I-65-412.20-2016 to H.C.R.), the German Research Foundation (DFG) (SFB1399 – grant no. 413326622 to H.C.R.; J.G, R.K.T., M.L.S., S.v.K., M.P., M.F., H.G., R.B., F.B.,J.Brägelmann, SFB1218 - grant no. 269925409 and CECAD EXC 2030 – grant no. 390661388 to M.B. and E.M.; SFB1451 - grant no. 431549029 to M.B.; Co 260/6-1 to A.A.H. and K-K.C.), the Else Kröner-Fresenius Stiftung (EKFS-2014-A06 to H.C.R., 2016_Kolleg.19 to H.C.R., R.D.J.), the Deutsche Krebshilfe (1117240 to H.C.R., 70113041 to H.C.R. and F.B. and the Mildred Scheel Nachwuchszentrum Grant 70113307 to F.B. and J.Brägelmann.), the Boehringer Ingelheim Stiftung (Exploration Grant to M.B.), the European Research Council (ERC-StG-2015, grant number 677844 to M.B.), as well as the German Ministry of Education and Research (BMBF e:Med Consortium InCa, grant 01ZX1901 and 01ZX2201A to R.K.T., J.G., H.C.R, S.v.K., M.L.S., M.P., J.Brüning.).

Additional Funding was received from the program “Netzwerke 2021”, an initiative of the Ministry of Culture and Science of the State of North Rhine-Westphalia for the CANTAR project to J.G., R.K.T. and M.P. The results shown here are in part based upon data generated by the TCGA Research Network: https://www.cancer.gov/tcga. The data used for the analyses described in this manuscript were obtained in part from the GTEx Portal on 28/08/2020

